# *APOE*^ε3/ε4^ and *APOE*^ε4/ε4^ genotypes drive unique gene signatures in the cortex of young mice

**DOI:** 10.1101/2020.10.28.359422

**Authors:** Kate E. Foley, Dylan T. Garceau, Kevin P. Kotredes, Gregory W. Carter, Michael Sasner, Gareth R. Howell

## Abstract

**Background:** Restrictions on mouse models have significantly impacted research towards understanding the most common genotype contributing to dementia in the human population – *APOE*^*ε3/ε4*^. To address this, as part of MODEL-AD, we created new versions of humanized *APOE*^*ε4*^ and *APOE*^*ε3*^ mice on a C57BL/6J background that allow for unrestricted distribution and breeding.

**Methods:** To determine similarities and differences between *APOE*^*ε3/ε4*^ and *APOE*^*ε4/ε4*^ risk genotypes, we analyzed peripheral lipid concentrations as well as performed unbiased transcriptional profiling of the cortex at two and four months of age, comparing *APOE*^*ε3/ε4*^ and *APOE*^*ε4/ε4*^ to the reference *APOE*^*ε3/ε3*^. To further compare APOE genotypes, cohorts of *APOE*^*ε3/ε3*^, *APOE*^*ε3/ε4*^, and *APOE*^*ε4/ε4*^ mice were exercised by voluntary running from 1 month to 4 months of age.

**Results:** Cholesterol composition was significantly influenced by APOE genotype as early as 2 months, while triglycerides were affected by APOE genotype at 4 months. Importantly, RNA-sequencing of the cortex followed by linear modeling or weighted gene co-expression network analysis (WGCNA) revealed that the *APOE*^*ε3/ε4*^ genotype showed unique transcriptomic signatures to that of *APOE*^*ε4/ε4*^. Functional enrichment of the *APOE*^*ε3/ε4*^, but not *APOE*^*ε3/ε4*^ genotype, revealed sulfur and heparin binding as significant terms at 2 months, and extracellular matrix and blood coagulation at 4 months. Further, cell specific contributions of significant genes identified endothelial cells as overrepresented in the *APOE*^*ε3/ε4*^ but not *APOE*^*ε4/ε4*^ genotype. WGCNA analysis confirmed findings from linear modeling but also predicted that running at a young age affects myelination and gliogenesis across APOE genotypes.

**Conclusions:** In summary, *APOE*^*ε3/ε4*^ genotype-specific effects were observed in cortical transcriptional profiles, suggesting therapies aimed at modifying APOE biology to treat dementias may need to be targeted to specific *APOE* genotypes.

## Background

Late Onset Alzheimer’s Disease (LOAD), characterized by amyloid plaques and tau tangles, and vascular dementia (VaD), characterized by cerebral strokes, are the top two most common forms of dementia. One allele of Apolipoprotein E (*APOE*), *APOE*^*ε4*^, has been identified as the greatest genetic risk factor for these dementias (1, 2). Two other APOE alleles are *APOE*^*ε3*^ and *APOE*^*ε2*^, which confer neutral or protective risk for dementia, respectively. Allele frequencies vary between cases and controls, with *APOE*^*ε2*^, *APOE*^*ε3*^, and *APOE*^*ε4*^ present at a rate of 8%, 78%, and 14%, respectively in unaffected subjects, and 4%, 59% and 37% in AD cases (3). The frequency of LOAD increases from 20% in a non-carrier of *APOE*^*ε4*^, to 47% when carrying one copy, and 91% when carrying two copies (4). In VaD, *APOE*^*ε4*^ predisposes individuals for increased risk for cerebrovascular disease and ischemic stroke, potentially up to 30%(5). Between *APOE*^*ε3*^ and *APOE*^*ε4*^, a single nucleic acid difference, changing a thymine (T) to a cytosine (C) at rs429358, results in an amino acid change from cysteine at position 112 to arginine (C112R) in the carboxy-terminal receptor region of the APOE protein. With *APOE*^*ε4*^ being less frequent than *APOE*^*ε3*^, the more common risk genotype in a given population is *APOE*^*ε3/ε4*^, with 41.1% of AD cases possessing the *APOE*^*ε3/ε4*^ genotype compared to 14.8% with *APOE*^*ε4/ε4*^.

Genetic variation between mouse *APOE* alleles has been proposed to modify phenotypes relevant to human aging and dementia (6). However, mouse strains do not carry equivalent *Apoe* alleles to those observed in humans potentially limiting the translatability of these studies. Therefore, humanized *APOE* mice have contributed to what we currently understand about the myriad of APOE mechanisms that may increase risk for dementia. *APOE*^*ε4*^ is predicted to increase risk for dementia through either gain of toxic function or loss of function which may be context dependent. APOE is a lipid trafficking protein, with *APOE*^*ε4*^ preferentially binding to VLDL, while *APOE*^*ε3*^ preferentially binds HDL (5). The effects of *APOE*^*ε4*^ are not limited to lipid and cholesterol homeostasis; research has shown that *APOE*^*ε4*^ affects neuronal health by negatively affecting neurogenesis and increasing neuronal toxicity, amyloid and tau protein aggregate formation through the involvement of seeding amyloid and the decreased ability to clear it from the brain, glucose homeostasis by competing with other APOE isoforms for the Low density lipoprotein receptor-related protein 1 (*LRP1*), and cerebrovascular health through tight junction and basement membrane degradation (7-9). Only more recently have some researchers started looking at the effects of APOE genotype at a young age, finding that there are structural and functional differences between genotypes in humans and mice (10-13).

While human and murine studies have parsed out important mechanisms by which the *APOE*^*ε4*^ allele differs in comparison to the *APOE*^ε3^ allele to increase detrimental brain pathology, little is known about the ways that *APOE*^*ε3*^ and *APOE*^*ε4*^ interact in those with the *APOE*^*ε3/ε4*^ genotype to affect risk for dementia. It is conceivable there may be compensatory, dominant, or *APOE*^*ε4*^ dose-dependent effects. This lack of knowledge has been due in part due to legal restrictions for the use of existing humanized *APOE* strains – including breeding of *APOE*^*ε4*^ to *APOE*^*ε3*^ mice to create *APOE*^*ε3/ε4*^ mice. Therefore, studies in mice have been limited to the *APOE*^*ε3/ε3*^ and *APOE*^*ε4/ε4*^ genotypes. In contrast, human studies are typically underpowered to separate *APOE*^*ε3/ε4*^ from *APOE*^*ε4/ε4*^, creating the much larger, ambiguous group of *APOE*^*ε4*^+ (or *APOE*^ε4^ carrier). To overcome these limitations, here we describe the creation of a new set of humanized *APOE* alleles, using a similar design to models widely used previously. We then use these humanized *APOE* mice to test our hypothesis that *APOE*^*ε3/ε4*^ mice show characteristics that may modify risk for dementias that are distinct from *APOE*^*ε4/ε4*^ mice.

## Methods

### Mouse Husbandry

All experiments involving mice were conducted with approval and accordance described in the Guide for the Care and Use of Laboratory Animals of the National Institutes of Health. All experiments were approved by the Animal Care and Use Committee at The Jackson Laboratory. Mice were kept in a 12/12-hour light/dark cycle and fed ad libitum 6% kcal fat standard mouse chow.

### Creation of humanized APOE mouse strains

Humanized APOE mice were created in collaboration with the Genetic Engineering Technologies core at The Jackson Laboratory. The mouse *Apoe* gene is located on chromosome 7 at 19,696,109 – 19,699,166. An *APOE*^*ε4*^ gene-targeting construct was made that included 4980 bp of mouse sequence, which defined the mouse 5’ homology arm including exon 1 of mouse *Apoe*, 4292 bp of human *APOE*^*ε4*^ sequence including human protein coding exons 2-4 of the human gene as well as an additional 1.5 kb of flanking human sequence after the 3’UTR to include any potential regulatory sequences. Exon 4 contained sequence that encoded the *APOE*^*ε4*^ isoform (nucleotide sequence for arginine at R130 and R176). Note that *APOE*^*ε4*^ includes an 18 amino acid signal peptide at the N-terminus of the protein such that *APOE*^*ε4*^ including the signal peptide is 317 amino acids and the variant amino acids which differ among *APOE*^*ε2*^, *APOE*^*ε3*^ and *APOE*^*ε4*^ are at positions 130 and 176. In the mature APOE proteins (299 amino acids), the variant amino acids which differ among *APOE*^*ε2*^, *APOE*^*ε3*^ and *APOE*^*ε4*^ are at positions 112 and 158. A Frt-neo-Frt (FNF) selection cassette was inserted after the human sequence followed by a Nde1 restriction site (for ease of Southern screening). The FNF cassette was followed by 5166 bp of mouse sequence, the 3’ homology arm. The resulting 14,438 bp synthesized construct was cloned into pBlight vector using recombineering techniques, producing a construct called mApoE_hAPOE4_PGKneo_mAPOE for gene targeting in embryonic stem cells. The *APOE*^*ε4*^ gene-targeting construct was introduced into cultured embryonic stem (ES) cells of a C57BL/6J (B6) mouse strain by electroporation. Homologous recombination produced loci that retained all normal mouse regulatory sequences (plus non-coding exon one) together with the human *APOE*^ε4^ protein-encoding exons 2-4. Transfected ES cells were screened by Southern blot to ensure correct targeting. Three clones were identified that were correctly targeted. ES cells containing the correctly targeted locus were introduced into albino B6 embryos, and the resultant chimeric mice were bred with B6 mice. Offspring carrying the modified locus in the germ-line were interbred to generate the homozygous genetically modified genome. All F1 matings produced normal litter sizes with a Mendelian distribution of the locus. Heterozygous animals were crossed to FLP recombinase expression mice (JAX Stock No. 005703) to remove the FRT site flanked PGK-neo cassette. Mice that no longer contained the FRT flanked PGK-neo cassette were then backcrossed to B6 at least once to remove the FLP recombinase transgene. SNP analysis was performed to validate the B6 background. Offspring that were negative for the FLP recombinase transgene were then interbred and maintained as *APOE*^*ε4/ε4*^ homozygous mice (JAX Stock No. 000664).

The *APOE*^*ε3*^ model was generated using CRISPR/Cas9-mediated gene targeting in *APOE*^*ε4*^ zygotes. An *APOE*^*ε4*^ specific sgRNA (GCGGACATGGAGGACGTGCG <u>CGG</u>; PAM site is underlined) was used to target the human knock-in allele just downstream of the valine codon that defines the *APOE*^*ε4*^ allele, and a 129nt single stranded oligonucleotide (GAGACGCGGGCACGGCTGTCCAAGGAGCTGCAGGCGGCGCAGGCCCGGCTGGGCGCGGACATGGAGGAC<u>GTc</u>**T**GCGGCCGCCTGGTGCAGTACCGCGGCGAGGTGCAGGCC ATGCTCGGCCA) including the CGC->**T**GC substitution (bold) to change arginine to cysteine and silent mutation GTG-GT<u>C</u> (underlined) to prevent re-cutting. Putative founders were bred to B6 mice for at least four generations and then interbred to be maintained as *APOE*^ε3/ε3^ homozygous mice (JAX Stock No. 029018).

To create experimental cohorts, *APOE*^*ε3/ε3*^ mice were crossed to *APOE*^*ε4/ε4*^ mice to generate *APOE*^*ε3/ε4*^ mice, which were then intercrossed to give littermate controls across all required genotypes (*APOE*^*ε3/ε3*^, *APOE*^*ε3/ε4*^, and *APOE*^*ε4/ε4*^).

### Genotyping of APOE mice

Genotyping to differentiate between *APOE*^*ε3*^ and *APOE*^*ε4*^ alleles was performed via ear punch at one month of age. A gel based PCR assay was used to determine insertion of the humanized construct. Primers included a common forward spanning over intron 1 and exon 2 (AATTTTTCCCTCCGCAGACT), a wild type (WT) mouse reverse spanning intron 2 (ACAGCTGCTCAGGGCTATTG), and a humanized reverse (AGGAGGTTGAGGTGAGGATG). A band for WT (wild type - mouse control) shows at 244bp, while the humanized insertion results in a 148bp band. To differentiate between the *APOE*^*ε3*^ and *APOE*^*ε4*^ alleles, sanger sequencing was used to identify CT for *APOE*^*ε3*^, and GC for *APOE*^*ε4*^, and a double peak signifying the presence of both *APOE*^*ε3*^ and *APOE*^*ε4*^ (GT).

### Validation of APOE expression by Western Blotting

Snap-frozen hemispheres were homogenized by hard tissue homogenizer (USA Scientific, Ocala, FL) and lysed in 700mL RIPA buffer (R0278, Sigma, St. Louis, MO) supplemented with 100X protease and phosphatase inhibitor reagents (1861281, Thermo Fisher Scientific, Waltham, MA). Lysates were incubated for 1 hour at 4°C before pelleting insoluble proteins by spinning at 4°C, 11,000xg for 15 minutes. Protein concentration was determined by Bradford protein assay (Biorad, Hercules, CA), according to manufacturer’s instructions. Samples were mixed with 10x Laemlli buffer (42556.01, Amsbio, Cambridge, MA), boiled for 10 minutes, and run on 12% SDS PAGE gels (456-1044, BioRad) with colorimetric ladder (RPN800E, GE, Boston, MA). Gels were transferred to PVDF membranes for immunoblotting and imaging using an iBlot2 dry blotting system (Thermo Fisher). Membranes were blocked in 5% non-fat dry milk in 1xPBS+0.1% Tween20 for 1 hour prior to incubating with primary antibodies diluted in 5% non-fat dry milk in 1xPBS+0.1% Tween20 for 1 hour at room temperature. Membranes were washed in 1xPBS+0.1% Tween20 before incubating with secondary antibodies diluted in 5% non-fat dry milk in 1xPBS+0.1% Tween20. HRP-conjugated secondary antibodies targeting primary antibody host IgG were incubated at 1 hour at room temperature. Membranes were washed in 1xPBS+0.1% Tween20 before digital imaging with SuperSignal West Pico PLUS chemiluminescent substrate (34579, Thermo Fisher). Antibodies used include: Pan human APOE-Millipore AB947; Mouse Apoe - Novus NB100-240; human APOE^ε4^ – Novus NBP1-49529; Actin – Abcam ab179467.

### Exercise by Voluntary Running

At first, mice were group housed (two or three per pen) and were provided access to low profile saucer wheels (Innovive Inc) 24 hours a day from 1 month to 4 months. Sedentary mice were not provided access to running wheels. At 4 months, just prior to harvest, mice were separated and individually housed with a trackable low-profile running wheel (Med Associates Inc.) or no wheel access. Rotations per minute during lights out (12 hours, 6:00pm – 6:00am) was quantified. Running wheel rotations were measured in 1-minute bins to allow for distance traveled (sum of rotations) calculated per mouse each night. Average rotations were calculated per mouse. Average speed while active was calculated by isolating the minute intervals where activity was measured (>0) and averaging the number of rotations for the minutes active. Percent of time at each speed was calculated by totaling the number of minute bins that mice ran between 0, 1-30 rotations, 31-70 rotations, 71-100 rotations and 100+ rotations and dividing by the total amount of minutes tracked. Any nights that had fewer than 700 minutes tracked were excluded from analysis.

### Harvesting, Tissue Preparation and Blood Lipid Profiling Assessment

All mice were euthanized by intraperitoneal injection of a lethal dose of Ketamine (100mg/ml)/Xylazine(20mg/ml) and blood was collected in K2 EDTA (1.0mg) microtainer tubes (BD) through approved cardiac puncture protocols. Mice were perfused intracardially with 1XPBS. Brains were carefully dissected and hemisected sagittally and the cortex (Ctx) was then carefully isolated and snap frozen in solid CO_2_ for RNA-sequencing. Blood was kept at room temperature for at least 30 minutes to prevent clotting and then centrifuged at 21°C for 10 minutes at 5000rpm. Plasma was carefully collected. Plasma lipid concentrations were characterized on the Beckman Coulter AU680 chemistry analyzer.

### RNA Extraction, Library Construction, RNA Sequencing, and RNA-seq quality control

RNA sequencing (RNA-seq) was performed by The Jackson Laboratory Genome Technologies Core. RNA extraction involved homogenization with TRIzol (Invitrogen) as previously described (14). RNA was isolated and purified using the QIAGEN miRNeasy mini extraction kit (QIAGEN) in accordance with manufacturer’s instructions. RNA quality was measured via the Bioanalyzer 2100 (Agilent Technologies) and poly(A) RNA-seq sequencing libraries were compiled by TruSeq RNA Sample preparation kit v2 (Illumina). Quantification was performed using qPCR (Kapa Biosystems). RNA-seq was performed on the HiSeq 4000 platform (Illumina) for 2×100bp reads for a total of 45 million reads according to the manufacturer’s instructions. Quality control for each sample was completed using NGSQCToolkit v2.3 which removed adaptors and trimmed low quality bases (Phred<30)(15). To quantify gene expression of the trimmed reads, we used RSEM v1.2.12 which uses Bowtie2 v2.2.0 for alignment of these reads (16). We used mouse genome mm-10 based upon the B6 reference genome.

### Linear Modeling (LM) and Functional Enrichment

Genes were filtered by 1) removing all genes that did not vary in expression and 2) removing all genes that did not have at least five reads in 50% of the samples. Remaining genes were normalized using DEseq2 (17). Principle component analysis (PCA) on the variance stabilized data (vst) identified 3 outliers from the 2 month dataset, and 5 outliers from the 4 month dataset which were excluded from further analysis. *APOE* genotype will be referred to as ‘genotype’. A linear model was used on the normalized counts to identify genes significantly fit by the predictors of our model: sex, genotype, and sex-genotype in the 2 month dataset, and sex, genotype, activity, and genotype-activity in the 4 month dataset. Reference data in the model were ‘female *APOE*^*ε3/ε3*^*’* for 2 month and ‘female sedentary *APOE*^*ε3/ε3*^’ for 4 month data. The linear model was performed on 17,763 genes for 2 months and on 18,987 genes for 4 months. Significant genes were identified per predictor term (sex, genotype, etc.) based upon Pr(>|t|<0.05). Functional enrichment for each linear model predictor term was performed using the clusterProfiler package with p<0.10 to determine significant gene ontology (GO) terms (18). Background lists consisted of all genes prior to significant geneset filtering.

### Cell type specific enrichment

Publicly available cell type gene expression was downloaded from the brainrna-seq.org dataset generated by the Barres laboratory (19, 20). Significant genes for each 4 month linear model predictor term were cross referenced to this dataset. Some genes in our datasets were not found in the brainRNA-seq dataset and were excluded. Number of genes per cell type was expressed as a percentage of a specific cell type divided by all genes that were able to be cross referenced for the linear model predictor terms.

### WGCNA

Weighted gene co-expression network analysis required the WGCNA package by Horvath and Langfelder (21, 22). Normalized and variance transformed data (vst) without outliers was used to build weight gene co-expression networks. First, all samples passed the function goodSamplesGenes to check for incomplete sample data. Samples were clustered to identify outliers using hierarchical clustering; two samples were excluded from further analysis. Next, the soft-thresholding power (β) was chosen by calculating scale-free topology through the relationship between power and scale independence which resulted in a softPower of 5. Then 18 modules were clustered based upon minModuleSize = 30, and a mergeCutHeight = 0.4. Module eigengenes for each module were identified and the correlation to sex, genotype, and activity was computed. The p values were corrected for multiple testing by using the false discovery rate. Genes in each module were annotated by using the AnnotationDbi package (23). Module membership (module eigengene correlated with gene expression) was regressed against gene significance (the correlation between the linear model predictor term and each individual gene) to identify highly associated modules. Genes in a module were processed through clusterProfiler to identify functional enrichment.

### Statistical Analysis

Statistical analysis for 2 month biometric data including cholesterol composition, triglyceride, unfasted glucose, and non-esterified fatty acids (NEFA) were calculated by two-way ANOVA (interactions tested: sex-genotype, main effects: sex, genotype), and a post-hoc Tukey test in GraphPad Prism v7.0a. Statistical analysis for 4 month biometric data was performed by three-way ANOVA (interactions tested: sex-genotype-activity, genotype-activity, sex-activity, sex-genotype, main effects tested: sex, activity, genotype) followed by a TukeyHSD post-hoc test in R v1.2.1335. Terms were considered significant if p<0.05. Post-hoc tests were used to determine differences between genotypes per sex, significant effects between genotypes and sexes were not reported (i.e. female-sed-*APOE*^*ε3/ε4*^ to male-sed-*APOE*^*ε3/ε3*^). All weight and biometric data included groups of 5-16 mice per sex/genotype/activity. Transcriptional profiling was performed on groups of 6 mice per sex/genotype/activity. Details of statistical analyses of linear modeling, functional enrichment, and WGCNA data are provided above.

## Results

### Generation and validation of humanized APOE^ε3/ε3^ and APOE^ε4/ε4^ mice

Due to breeding restrictions on commonly used humanized *APOE* models, *APOE*^*ε3/ε3*^ and *APOE*^*ε4/ε4*^ mice were created on the well characterized B6 mouse strain using a similar design to that described previously (**See methods**) (24). Briefly, exons two, three, and four of the mouse *Apoe* gene on chromosome 7 were replaced by exons two, three and four of human *APOE* through homologous directed repair (HDR) (**Fig 1A)**. Exon 4 of the *APOE* gene sequence encoded the *APOE*^*ε4*^ risk allele incorporating cytosines at rs7412 and rs428358 which result in arginine at amino acid positions 130 and 176. To confirm complete target replacement in *APOE*^*ε4/ε4*^ mice, we used a PCR based assay combined with Sanger sequencing (**Fig 1B, C)**. *APOE*^*ε3/ε3*^ mice were then produced using CRISPR-Cas9 of *APOE*^*ε4/ε4*^ mice to replace cytosine with thymine at rs429358, which results in a cysteine at amino acid position 130, creating the *APOE*^*ε3*^ allele. The *APOE*^*ε3/ε3*^ strain was also validated using PCR and Sanger sequencing (**Fig 1B, C)**. Presence of human APOE protein was assessed in each strain by western blot using a pan anti-human APOE antibody along with an anti-APOE^ε4^ antibody, showing that human APOE is present in both the humanized APOE mice, but APOE^ε4^ was only detectable in *APOE*^*ε4*^ mice (**Fig 1D, Supp Fig 1)**.

**Figure 1:**
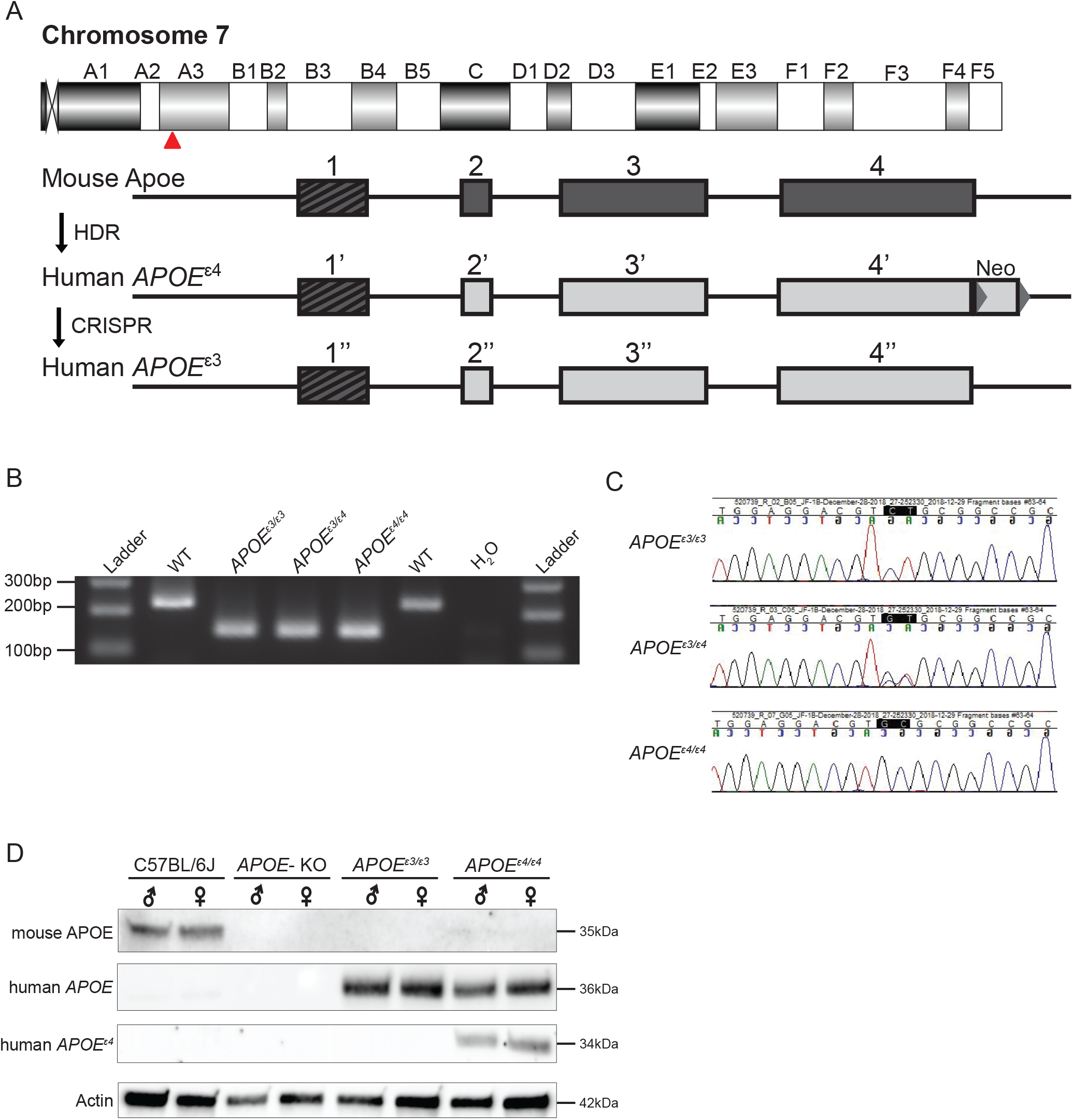
Creation and validation of *APOE* allelic series. **A**. Schematic of human *APOE* allele insertion at the mouse *Apoe* locus on chromosome 7. Exon 1 remained unchanged (mouse) and exons 2-4 were replaced with human *APOE*^*ε4*^ sequence through homology directed repair (HDR). The *APOE*^*ε3*^ SNP was changed through CRISPR mediated endonuclease activity. Critical amino acids differing between *APOE*^*ε4*^ and *APOE*^*ε3*^ noted in red. **B**. PCR based assay to identify humanized *APOE* insertions vs WT mouse *Apoe*. **C**. Sanger sequencing to distinguish *APOE*^*ε3*^ and/or *APOE*^*ε4*^ alleles. **D**. Western blot of mouse *Apoe*, pan human *APOE*, and human *APOE*^*ε4*^-specific expression.

### Biometrics show cholesterol composition differs by sex and APOE genotype at 2 months of age

To begin to evaluate this new humanized *APOE* strain, *APOE*^*ε3/ε3*^ mice were crossed to *APOE*^*ε4/ε4*^ mice to create *APOE*^*ε3/ε4*^ mice that were then intercrossed to create cohorts of male and female mice carrying all APOE genotypes (*ε3/ε3, ε3/ε4, ε4/ε4*). Circulating APOE protein has many functions, including lipid trafficking and mediating homeostatic levels of cholesterol. Therefore, to determine the effects of *APOE* genotype on biometric measurements at 2 months, first weight across genotypes and sex was assessed. Weight was significantly different between sexes, but not between genotypes, with males weighing more than females (**Fig 2A**). Next, plasma lipid concentrations were determined and compared across sex and *APOE* genotype using an ordinary two-way ANOVA. Both total cholesterol and low-density lipoprotein (LDL) concentrations showed a significant interaction between sex and *APOE* genotype. Posthoc tests revealed that total cholesterol concentrations were significantly higher in male compared to female *APOE*^*ε4/ε4*^ mice (p<0.05, **Fig 2B)**. LDL concentrations were significantly higher in female *APOE*^*ε3/ε3*^ mice compared to both male *APOE*^*ε3/ε3*^ (p<0.001) and female *APOE*^*ε4/ε4*^ mice (p<0.01, **Fig 2C**). High-density lipoprotein (HDL) concentrations were significant for sex, with HDL levels higher in males than females across all *APOE* genotypes (**Fig 2D**). Triglyceride concentrations have been previously reported to be higher in those with the *APOE*^*ε4/ε4*^ over the *APOE*^*ε3/ε3*^ genotype, but at 2 months there was no significant difference between the *APOE* genotypes (**Fig 2E**). Unfasted glucose concentrations were significantly affected by sex at this 2-month timepoint, with increased glucose concentrations in males compared to females (**Fig 2F**). These data show that *APOE* genotype influences cholesterol composition in mice as early as 2 months. However, posthoc tests showed no significant differences comparing the *APOE*^*ε3/ε4*^ genotype to either *APOE*^*ε3/ε3*^ or *APOE*^*ε4/ε4*^.

**Figure 2:**
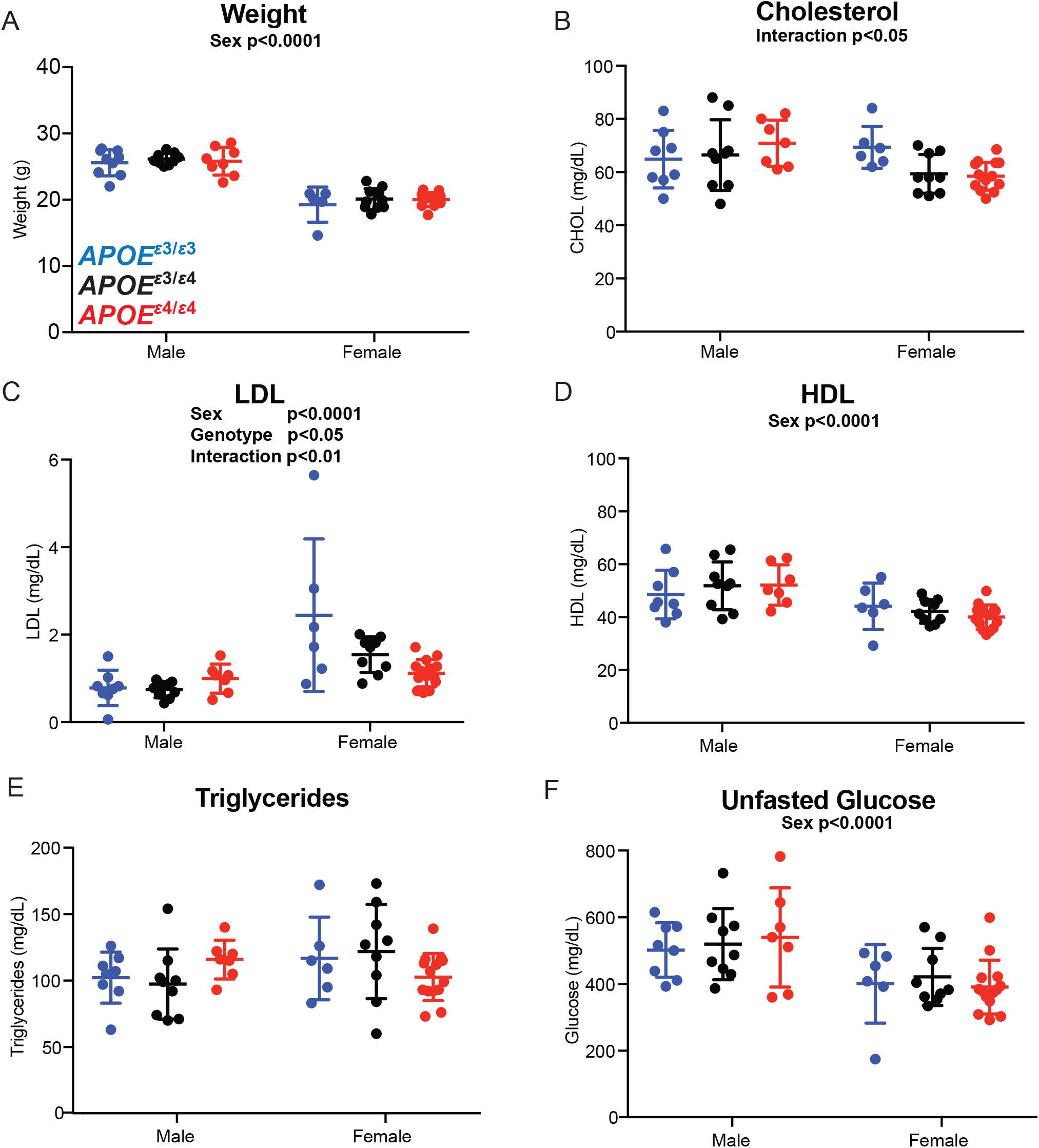
Plasma cholesterol composition affected by both *APOE* genotype and sex at 2 months. **A**. Weight (grams. g) at harvest in both male and female *APOE* mice showed expected significant sex difference. **B**. Total plasma cholesterol concentration revealed a significant interactive effect of *APOE* genotype and sex. **C**. Low density lipoprotein (LDL) concentration showed a significant interactive effect between *APOE* genotype and sex. **D**. High density lipoprotein concentration (HDL) showed a significant sex effect. **E**. Triglyceride concentration showed no effect by *APOE* genotype or sex. **F**. Unfasted glucose showed significant sex effect.

### Transcriptional profiling of brains of young APOE mice show unique enrichment for APOE^ε3/ε4^

Mathematical modeling and gene set enrichment are being widely used in human genomics to define molecular processes relevant to AD and dementia (25-27). Therefore, we adopted this strategy to evaluate differences between one or two *APOE*^*ε4*^ alleles compared to the reference (neutral) *APOE*^*ε3/ε3*^ genotype in our mouse models. To compare the brain transcriptomes of each of the three *APOE* genotypes, RNA-seq was performed on cortical tissue collected from 2-month-old male and female littermate controls. Principle component analysis (PCA) showed separation between the sexes, however the *APOE* genotypes did not differentiate at this young age (**Fig 3A,B**). This result was not unexpected as previous studies also did not see a clear distinction between *APOE* genotypes in 3-month-old mice by PCA (10). Linear modeling was then used to identify genes that were expressed as a function of sex, *APOE* genotype, and the interaction between sex and genotype (sex-genotype interaction) (**Fig 3C, see Methods**). This model used pairwise comparisons to look at genotype effects; separately comparing *APOE*^*ε3/ε4*^ or *APOE*^*ε4/ε4*^ genotypes to *APOE*^*ε3/ε3*^, to evaluate the significant linear relationships between one or two *APOE*^*ε4*^ alleles compared to reference (p<0.05). Results showed 456 genes significant for the factor sex-*APOE*^*ε4/ε4*^, 139 genes significant for sex-*APOE*^*ε3/ε4*^, 423 genes significant for sex, 284 genes significant for *APOE*^*ε4/ε4*^, and 196 genes significant for *APOE*^*ε3/ε4*^. There were more genes affected by the interaction of sex and *APOE*^*ε4/ε4*^ (456 significant genes, p<0.05), than the interaction between sex and *APOE*^*ε3/ε4*^ (139 significant genes, p<0.05) (**Fig 3C**). These results suggest an interaction between sex and *APOE* genotype even at this young age, which may contribute to the increased risk for cognitive decline and dementia in *APOE*^ε4^ carrier females compared to males later in life. GO term enrichment on these gene sets was performed using clusterProfiler (18). No GO terms were enriched for the genes significant for the sex-*APOE*^*ε3/ε4*^ interaction or the sex-*APOE*^*ε4/ε4*^ interaction, suggesting multiple unrelated processes may be subtly affected, and may represent early biological disruptions that increase with age.

**Figure 3:**
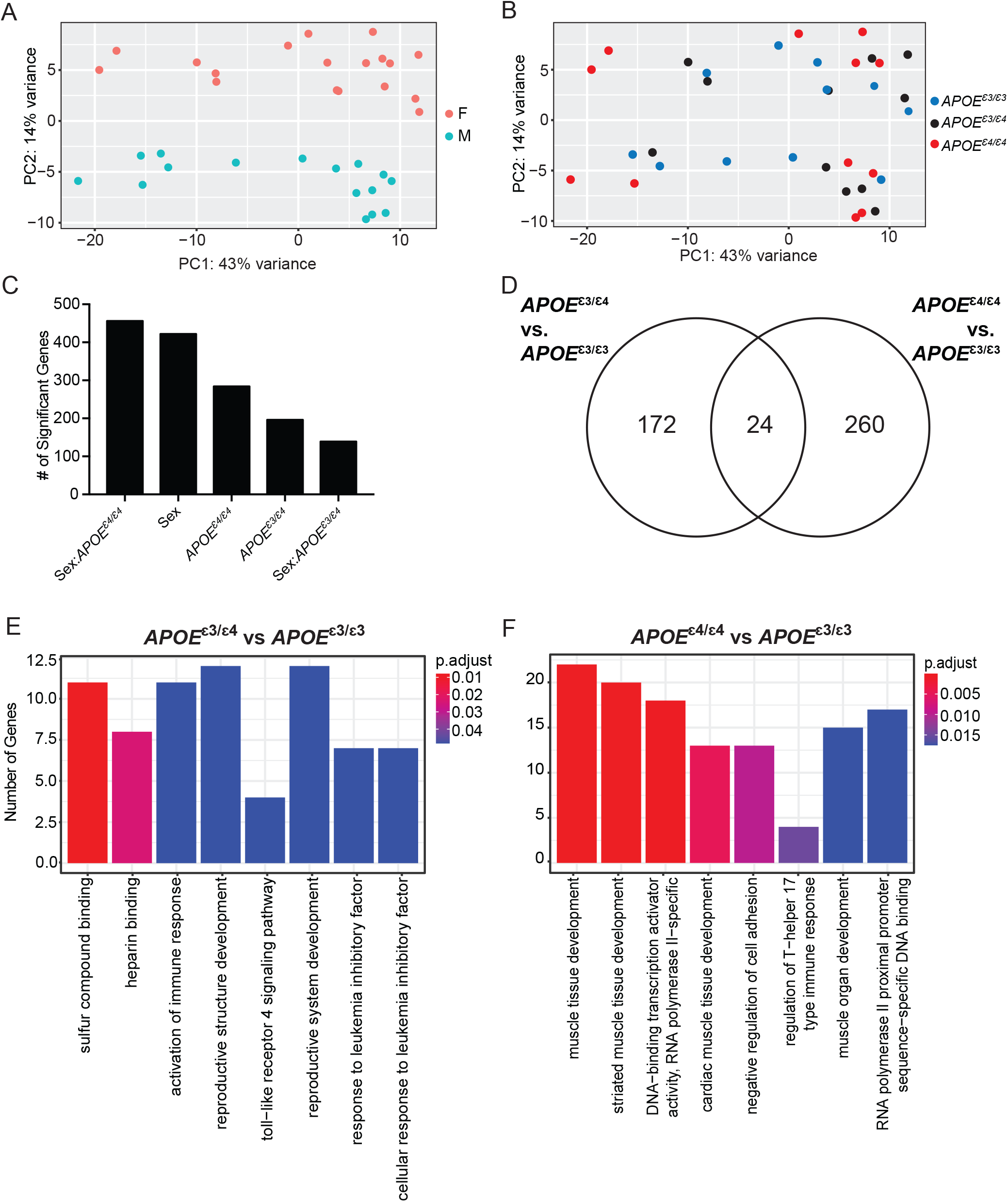
Cortical transcriptome analysis revealed differential functional enrichment between *APOE*^*ε3/ε4*^ and *APOE*^*ε4/ε4*^ genotype at 2 months. **A**. PCA showed distinct clusters of male and female samples. **B**. However, PCA showed samples do not cluster by *APOE* genotype at 2 months. **C**. Number of significant genes for each linear model predictor term. **D**. Number of significant genes unique to 2 months *APOE*^*ε4/ε4*^ when compared to *APOE*^*ε3/ε3*^, and 2 month *APOE*^*ε4/ε4*^ compared to *APOE*^*ε3/ε3*^, as well as the number of significant genes intersecting both terms. **E**. Functional enrichment of the 172 significant genes unique to *APOE*^*ε3/ε4*^ compared to *APOE*^*ε3/ε3*^. **F**. Functional enrichment of the 260 significant genes unique to *APOE*^*ε4/ε4*^ compared to *APOE*^*ε3/ε3*^.

To determine the functional relevance of the transcriptional differences in *APOE*^*ε3/ε4*^ and *APOE*^*ε4/ε4*^ mice compared to *APOE*^*ε3/ε3*^, independent of sex, we focused on the 196 genes significant for *APOE*^*ε3/ε4*^, and the 284 genes significant for *APOE*^*ε4/ε4*^. Hypothesizing that there was a simple additive effect of *APOE*^*ε4*^ allele dosage on gene expression, we expected that many of the genes in these two lists would overlap, with a few genes unique to *APOE*^*ε3/ε4*^ alone. However, only 24 genes (12% of the *APOE*^*ε3/ε4*^ genes, 8% of the *APOE*^*ε4/ε4*^ genes) intersected, suggesting different biological processes are modified in *APOE*^*ε3/ε4*^ or *APOE*^*ε4/ε4*^ when compared to *APOE*^*ε3/ε3*^ mice (**Fig 3D**). Enrichment of the significant genes for *APOE*^*ε3/ε4*^ identified GO terms such as ‘sulfur compound binding,’ ‘heparin binding,’ and ‘activation of immune response’ (**Fig 3E**). Genes associated with these GO terms included *Prelp, Vtn*, and *C2* (**Supp Fig 2A**). *Prelp* and *Vtn* contribute to ‘heparin binding’ by acting as critical matrix anchoring, and cell adhesion molecules. Complement component C2 - a key component of the classical complement pathway – is associated with ‘activation of immune response’ (28). In contrast, GO terms that were enriched for *APOE*^ε4/ε4^ when compared to *APOE*^*ε3/ε3*^, identified ‘muscle tissue development’, ‘cardiac muscle tissue development’, and ‘negative regulation of cell adhesion’ (**Fig 2F**). Genes with these GO terms include *Gli1, Klf4*, and *Sox8* (**Supp Fig 2B**). These three transcription factors are associated with cell proliferation, cell cycle, and cell fate respectively (29-34). Based on our linear modeling using *APOE* ^*ε3/ε3*^ as the reference strain, these results demonstrate that in mice as young as 2 months, cortical gene expression signatures appear differentially affected by the number of *APOE*^*ε4*^ alleles present, with the *APOE*^*ε3/ε4*^ genotype modulating genes relevant to immune, heparin, and sulfur processes, as opposed to the *APOE*^*ε4/ε4*^ genotype that modulated genes involved in muscle development and cell adhesion.

### Biometrics reveal sex and activity, but not APOE genotype as a major factor in plasma chemistry

Physical activity, especially running, has been widely considered a beneficial intervention to reduce risk for age-dependent cognitive decline and dementia. It is unknown, however, whether running has a similar beneficial effect across *APOE* genotypes – particularly when comparing *APOE*^*ε3/ε4*^ to *APOE*^*ε4/ε4*^. To evaluate whether running has differential effects across *APOE* genotypes at a young age, male and female *APOE*^*ε3/ε3*^, *APOE*^*ε3/ε4*^ and *APOE*^*ε4/ε4*^ mice were given access to a voluntary running wheel for 12 weeks from 1 months to 4 months (**Fig 4A**). Sedentary littermate controls were not given access to a running wheel. Running was tracked during the last week of the experiment (**see Methods**). Of the mice that ran, there was some expected variation between mice for amount of time spent running (35) (**Supp Fig 3A,B**). Number of rotations were averaged and showed no significant differences between *APOE* genotypes for both sexes (**Supp Fig 3C,D**). As has been shown before, female mice tended to run farther than male mice (35). Running speed was calculated and showed no observable differences between *APOE* genotypes for both sexes (**Supp Fig 3E,F, see Methods**). Weights were evaluated throughout the experiment, as well at harvest (4 months) and were compared across sexes and genotypes (**Fig. 4B-D**). There was a significant difference in weight between sexes at 4 months, however there were no differences in weight between the *APOE* genotypes for either sex, or between running and sedentary mice between sexes. The lack of weight differences between running and sedentary mice is likely due to the young age and short nature of the running paradigm. Unlike at 2 months which showed *APOE* genotype differences in cholesterol composition, *APOE* genotype was not significant for cholesterol composition at 4 months (**Fig 4E-G**). However, there was a significant sex-activity interaction effect for both total cholesterol as well as HDL concentrations (**Fig 4E,G**). Posthoc tests revealed that male sedentary cholesterol concentrations were significantly greater than female sedentary cholesterol concentrations (p<0.0001, **Fig 4E**). For HDL, in males there was a decrease in HDL concentration due to running (p<0.05), while in females there was a trending increase in HDL concentration with running (p=0.06), with the largest significant difference between male and female sedentary HDL concentrations (p< 0.0001, **Fig 4G**). Females showed a significantly greater LDL concentration than males (p<0.005, **Fig 4F**). Additionally, there was a significant sex-activity-genotype interaction effect in triglyceride concentration, which was unexpected because at 2 months there was no effect of sex or genotype (**Fig 4H**). There was a sex effect for unfasted glucose at 4 months, with glucose concentrations significantly higher in males than females (p< 0.0001, **Fig 4I**). There was no significant effect of sex, genotype or activity on NEFA (**Fig 4J**). Together, based on the assays performed, these results indicate that the main difference in lipid biochemistry at 4 months is driven by sex, while sex drove activity-dependent differences. Interestingly, *APOE* genotype did not have a significant effect on lipid biochemistry concentrations at this 4-month timepoint.

**Figure 4:**
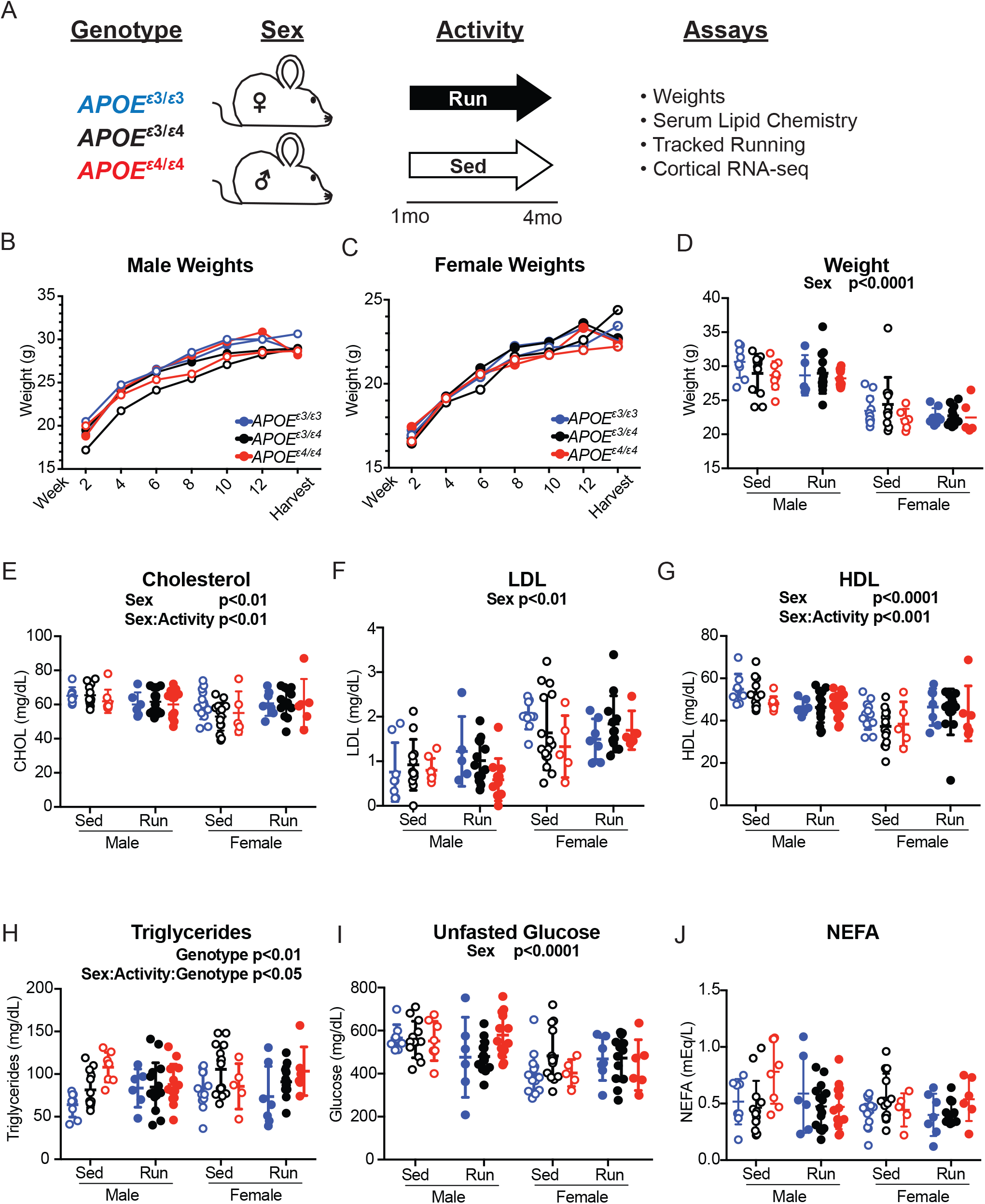
*APOE* genotype did not affect weight or cholesterol composition at 4 months. **A**. Schematic of experiment and timeline. **B and C**. Weights of male (**B)** and female (**C**) mice over the course of the running experiment. **D**. Weight at harvest of both male and female mice across *APOE* genotypes showed a significant effect of sex. **E**. Plasma total cholesterol concentrations revealed a main effect of sex and an interactive effect between sex and activity. **F**. LDL concentration revealed a significant effect of sex. **G**. HDL concentration revealed a significant effect of sex and interactive effect between sex and activity. **H**. Triglyceride concentrations revealed a significant effect of *APOE* genotype and an interactive effect between sex, activity and *APOE* genotype. **I**. Unfasted glucose concentrations revealed an effect of sex. **J**. NEFA concentrations revealed no significant effects.

### Genes significant for APOE^ε3/ε4^ genotype enrich for ECM- and coagulation-related terms

Transcriptional profiling was performed on cortical tissue isolated from six mice per sex/genotype/activity. Mice were selected across a range of distances ran (**Fig 5A, B and Supp Fig 3C, D**) based on previous experiences that mice show natural variation in voluntary running (35). We used a similar linear modeling approach as previously stated (comparing either the *APOE*^*ε3/ε4*^ or *APOE*^*ε4/ε4*^ genotypes to reference *APOE*^*ε3/ε3*^ genotype) to evaluate gene expression as a function of sex, *APOE* genotype, activity, and the interaction between genotype and activity (genotype-activity interaction).. Results showed 1,102 genes significant for sex, 540 genes significant for activity, 488 genes significant for *APOE*^ε4/ε4^, 405 genes significant for *APOE*^ε4/ε4^-activity interaction, 327 genes significant for *APOE*^ε3/ε4^, and 289 genes significant for *APOE*^ε3/ε4^-activity interaction (p<0.05, **Fig 5C**). The largest number of genes were significant for sex which enriched for ‘response to interferon beta’, ‘response to virus’, and ‘innate immune response’, suggesting a difference in immune function between the sexes (**Supp Fig 4**). *Ifitm3, Clu*, and *Plgc2* were among the genes significant for sex. Interferon-induced transmembrane protein 3 (*Ifitm3*) is a critical interferon response gene that increases in expression with AD and was recently reported as a modifier of gamma secretase activity (36). Clusterin (*Clu*), functions as a chaperone protein and regulates immune- and lipid-related functions (36). A common variant in *Clu* (up to 88% prevalence) increases risk for AD. Phospholipase C gamma 2 *(Plcg2*) is also commonly associated with AD, shows higher expression in AD than controls, and functions as a cell surface signal relay protein (37).

**Figure 5:**
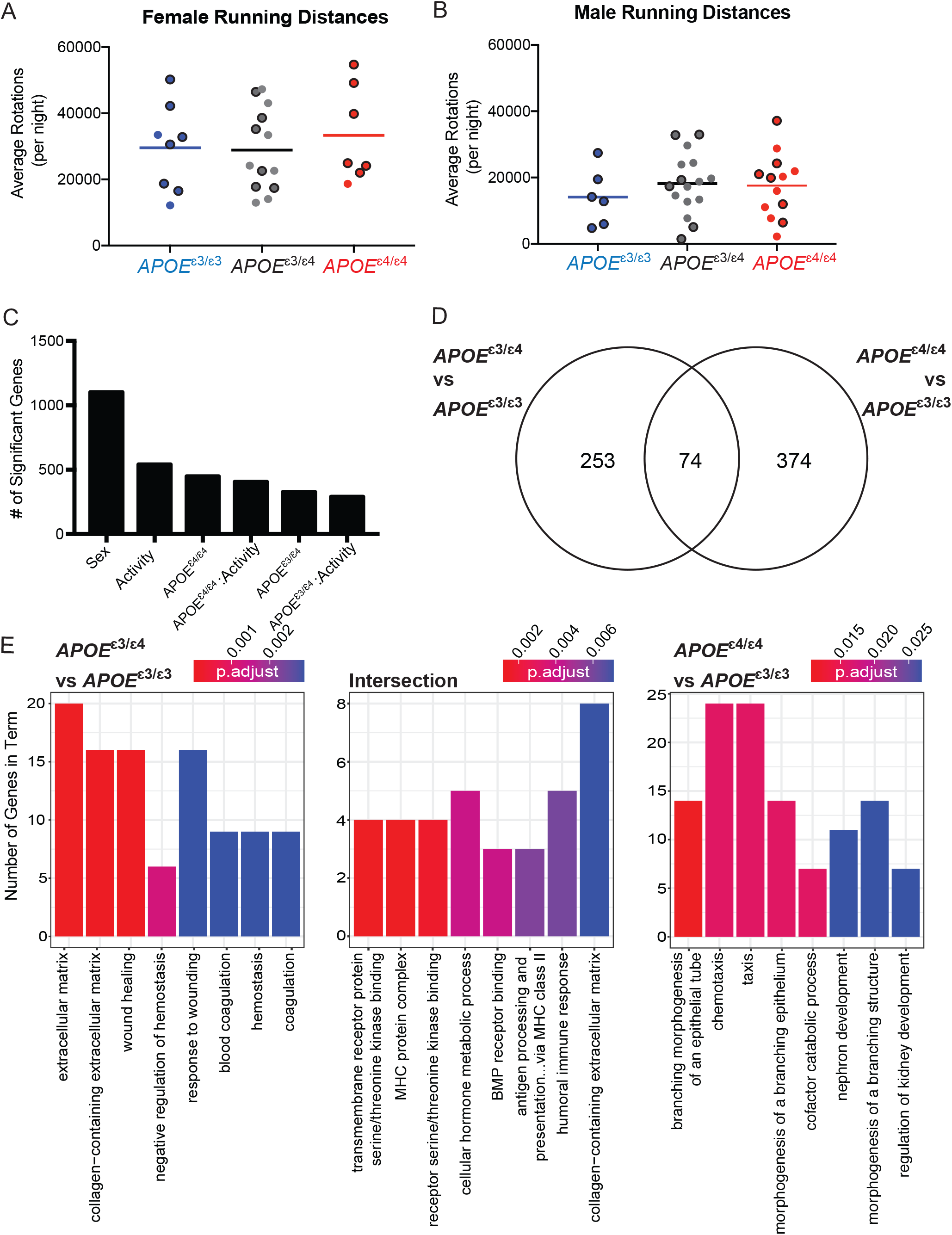
Significant genes unique for *APOE*^*ε3/ε4*^ genotype showed functional enrichment for ‘extracellular matrix’ and ‘response to wounding’ at 4 months. **A and B**. Average number of rotations per night during the dark cycle for female (**A**) and males (**B**) across all *APOE* genotypes for the six mice per group that were selected for RNA-seq. Mean is representative of all mice, including those that were not sequenced (see also **Fig. S2**). **C**. Number of significant gens per each linear model predictor term **D**. Number of significant genes unique to *APOE*^*ε3/ε4*^ when compared to *APOE*^*ε3/ε3*^, and *APOE*^*ε4/ε4*^ compared to *APOE*^*ε3/ε3*^, as well as the number of significant genes intersecting both groups. **E**. Functional enrichment of the 253 genes unique to *APOE*^*ε3/ε4*^ compared to *APOE*^*ε3/ε3*^ (left), 374 genes unique to *APOE*^*ε4/ε4*^ compared to *APOE*^*ε3/ε3*^ (right) and intersection between the two groups (middle).

To assess functional relevance, we focused on the 327 genes significant for *APOE*^*ε3/ε4*^ and the 448 genes significant for *APOE*^*ε4/ε4*^, when compared to *APOE*^*ε3/ε3*^ (**Fig 5D**). From these gene lists, only 74 genes were present in both lists (intersection, 22% from *APOE*^*ε3/ε4*^ genes, 17% from *APOE*^*ε4/ε4*^ genes) (**Fig 5D**). Functional enrichment of the genes unique for *APOE*^*ε3/ε4*^ when compared to *APOE*^*ε3/ε3*^ (253 unique genes) revealed GO terms such as ‘extracellular matrix’, ‘collagen-containing extracellular matrix’, and ‘wound healing’ (**Fig 5E**). These terms predict cerebrovascular changes in *APOE*^*ε3/ε4*^ but not *APOE*^*ε3/ε3*^ genotype. We also interrogated which genes contributed to these functional enrichment terms. Significant genes for *APOE*^*ε3/ε4*^ compared to *APOE*^*ε3/ε3*^, included ‘*Serpinh1’*, ‘*Col1a2’*, and ‘*Mmp19’* (**Supp Fig 6A**). Next, for the 374 genes that were unique for *APOE*^*ε4/ε4*^ genotype compared to reference *APOE*^*ε3/ε3*^, functional enrichment identified ‘branching morphogenesis of an epithelial tube’, ‘chemotaxis’, and ‘taxis’ (**Fig 5C)**. Significant genes for *APOE*^*ε4/ε4*^ included ‘*Gli3’*, ‘*Cx3cr1’*, and ‘*Ccr1’* (**Supp Fig 5A**). These terms predict that the *APOE*^*ε4/ε4*^ genotype influences growth and movement of tube-like structures in the cortex. Finally, for the 74 genes common to both (1) *APOE*^*ε3/ε4*^ compared with *APOE*^*ε3/ε3*^ and (2) *APOE*^*ε4/ε4*^ compared with *APOE*^*ε3/ε3*^, enrichment analysis identified terms that included ‘transmembrane receptor protein serine/threonine kinase binding’, ‘MHC protein complex’, and ‘BMP receptor binding’ (**Fig 5C**). Genes associated with these GO terms included ‘*Bmp5’*, ‘*Bmp6’*, and ‘*Spp1*’ (**Supp Fig 5B**). While it may be assumed that the *APOE*^*ε3/ε4*^ genotype is an intermediate between *APOE*^*ε3/ε3*^ and *APOE*^*ε4/ε4*^, these data support a model where some of the effects driven by the *APOE*^*ε3/ε4*^ genotype are unique and highlight the importance of studying this AD risk genotype separately to the *APOE*^*ε4/ε4*^ genotype.

**Figure 6:**
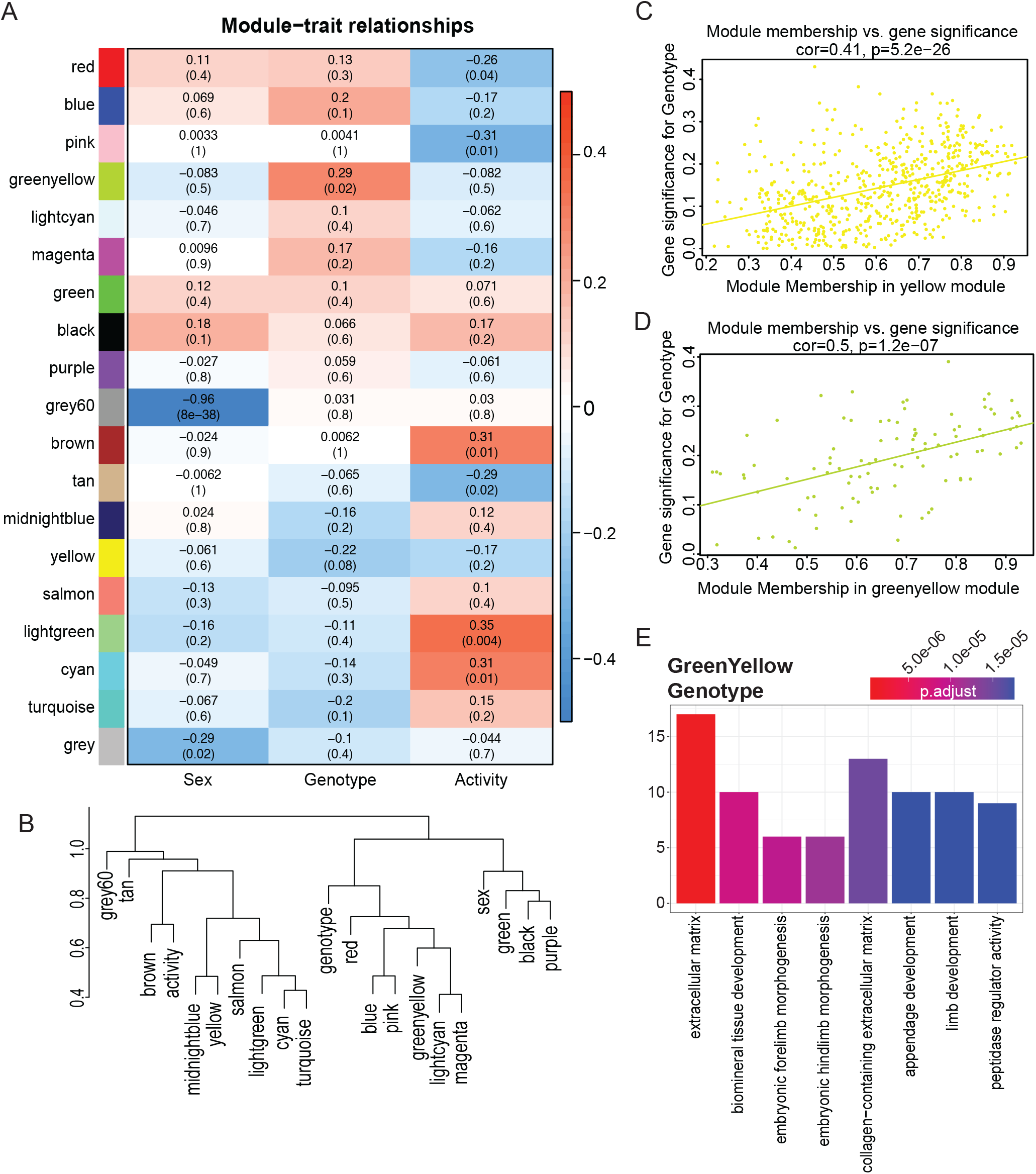
WGCNA confirmed functional enrichment of GO terms relating to extracellular matrix and morphogenesis is affected by *APOE* genotype. **A**. Heatmap of module correlation and FDR values for sex, *APOE* genotype, and activity. **B**. Eigengene dendrogram of each module and sex, *APOE* genotype and activity. Y axis is a measure of dissimilarity. **C and D**. Gene significance regressed against module membership to show correlation and significance for the yellow (**C**) and greenyellow (**D**) modules for *APOE* genotype. **E**. Functional enrichment of the genes in the greenyellow module which is significant for *APOE* genotype.

To determine cell types that may be responsible for the functional enrichment terms that were different between the *APOE* genotypes (*APOE*^*ε3/ε4*^ genotype compared to reference *APOE*^*ε3/ε3*^ and *APOE*^*ε4/ε4*^ genotype compared to reference *APOE*^*ε3/ε3*^), we cross-referenced the significant unique and intersection genes with a Brain RNA-seq dataset (19, 20). While many cell types can contribute to the ECM- and coagulation-related pathways, endothelial cells contributed to 33% of the unique genes that were significant for *APOE*^*ε3/ε4*^ (**Supp Fig 6B**). The cell types enriched for intersection-related terms included both oligodendrocyte precursor cells, as well as microglia (**Supp Fig 6B**). No specific cell types appeared to be responsible for branching morphogenesis- and taxis-related terms enriched in *APOE*^*ε4/ε4*^ unique terms, with a similar contribution by oligodendrocyte precursor cells, neurons, microglia, endothelial cells, and astrocytes. This suggests that while there is no single cell type contributing to the *APOE*^*ε4/ε4*^-dependent transcriptional differences, endothelial cells may be critical drivers of transcriptional differences in *APOE*^*ε3/ε4*^ mice. (**Supp Fig 6B**). For instance, transcriptional differences comparing *APOE*^*ε3/ε4*^ or *APOE*^*ε4/ε4*^ to *APOE*^*ε3/ε3*^ showed endothelial-related annexin genes including annexin 1 (*Anxa1*) and annexin 3 (*Anxa3*) had greater expression in *APOE*^*ε3/ε4*^, but not *APOE*^*ε4/ε4*^, when compared to *APOE*^*ε3/ε3*^ (**Supp Fig 6C**). Annexin 2 (*Anxa2*) showed a similar expression pattern in males, but in females there appears to be a stepwise increase with each *APOE*^*ε4*^ allele present (**Supp Fig 6C**). Matrix metalloprotease 25 (*Mmp25)*, an ECM relevant gene expressed by endothelial cells, also showed an *APOE*^*ε3/ε4*^-specific decrease in expression when compared to *APOE*^*ε3/ε3*^ and *APOE*^*ε4/ε4*^ (**Supp Fig 6C**). These results provide evidence that the *APOE*^*ε3/ε4*^ genotype, but not *APOE*^*ε4/ε4*^ genotype, is predicted to affect pathways involved in immune and cerebrovascular homeostasis.

To observe the effects of activity on mice carrying one or two *APOE*^*ε4*^ alleles, we next compared the genes for activity-*APOE*^*ε3/ε4*^ with activity-*APOE*^*ε4/ε4*^. Of the 289 genes significant for activity-*APOE*^*ε3/ε4*^ and the 405 genes significant for activity-*APOE*^ε4/ε4^, only 50 of these genes overlapped (intersection, 17% of activity-*APOE*^*ε3/ε4*^, 12% of activity-*APOE*^*ε4/ε4*^) (**Supp Fig 7A**). The 50 intersecting genes enriched for ‘anoikis’ (consisting of *Src, Tfdp1* and *Bmf*), suggesting a common cell death response between genotypes in response to activity (**Supp Fig 7B**,**C**). There was no significant enrichment for the genes unique to each interaction term. However, despite the lack of functional enrichment, these data suggest activity exerted differential effects on cortical gene expression in mice carrying one or two copies of *APOE*^*ε4*^.

#### WGCNA identifies myelination- and skeletal muscle cell differentiation-related terms associated with Activity

Linear modeling supported our hypothesis that one *APOE*^*ε4*^ allele in the *APOE*^*ε3/ε4*^ genotype has unique and different effects on transcriptional profiles than the presence of two *APOE*^*ε4*^ alleles (in *APOE*^*ε4/ε4*^), with respect to the reference *APOE*^*ε3/ε3*^. To further support our hypothesis, Weighted Gene Co-expression Network Analysis (WGCNA) was employed on the same 4-month dataset to create modules of similar expression patterns which can predict co-functional genes. WGCNA groups genes by correlated expression across the entire dataset, thus reducing complexing of the large number of input genes and producing biologically relevant gene modules. Nineteen modules were identified, and correlation and p-values were calculated for sex, *APOE* genotype, and activity (**Fig 6A**) with a p-value <0.1 considered significant. Two modules were significantly correlated with sex (grey60, grey), two modules significantly correlated for genotype (yellow and greenyellow), and six modules significantly correlated for activity (cyan, lightgreen, tan, brown, pink, and red) (**Fig 6A**). No modules were significant across multiple terms. Module eigengenes were calculated and dissimilarity plotted to visualize the relationship between modules and terms (**Fig 6B**). To evaluate gene contribution to each module, module membership and gene significance was computed. Module membership is the correlation of each individual gene in the module to the module expression (0-1). Gene significance for a trait is the association of each individual gene to the trait of interest. These measurements can be regressed, creating a correlation and p-value for the module membership and gene significance for the genes in the module. Both the yellow and greenyellow modules were significantly correlated to *APOE* genotype (yellow cor=0.41, greenyellow cor= 0.5) (**Fig 6C,D**). The genes comprising the yellow module were not significantly enriched for any GO terms. However, the genes that comprised the greenyellow module were enriched for GO terms that included ‘extracellular matrix’, ‘embryonic morphogenesis’, and ‘collagen-containing extracellular matrix’ (**Fig 6E**). The cyan and tan modules were significantly correlated for activity (cyan cor = 0.4, p=0.00026, tan cor = 0.26, p=0.0099) (**Supp Fig 8A**,**B**). Enrichment of the 79 genes in the cyan module revealed terms such as ‘ensheathment of neurons’, ‘myelination’, and ‘gliogenesis’ (**Supp Fig 8C**). Enrichment for the 84 genes in the tan module revealed terms such as ‘skeletal muscle cell differentiation’, ‘skeletal muscle organ development’, and ‘muscle organ development’ (**Supp Fig 8D**). Together, these results support findings from our previous analysis, which suggest that *APOE* genotype affects the extracellular matrix, collagen, and morphogenesis (**Fig 4)**. However, while there were no modules significant for *APOE* genotype and activity, an activity-dependent effect was observed for myelination, gliogenesis, and muscle development. These results show that running activity at a young age is predicted to affect myelination and gliogenesis processes across all APOE genotypes.

## Discussion

In this study we demonstrate the importance of including all human-relevant genotypes in our understanding of APOE biology. *APOE*^*ε3/ε4*^ is more common than *APOE*^*ε4/ε4*^ across Caucasian, African America, Hispanic, and Japanese populations, which means there is a stark disconnect between genotypes evaluated in mouse models to those present in human studies (3). In human studies, it is difficult to recruit enough *APOE*^*ε4/ε4*^ homozygous individuals which commonly results in a clustering of both *APOE*^*ε3/ε4*^ and *APOE*^*ε4/ε4*^. These strategies cloud differences between the two genotypes. In contrast, the vast majority of mouse studies performed to date primarily contrasted the *APOE*^*ε4/ε4*^ genotype with the *APOE*^*ε3/ε3*^ genotype. Our study here predicts the *APOE*^*ε3/ε4*^ genotype exerts unique effects both in the periphery and the brain suggesting more attention should be given to understanding the mechanisms by which the *APOE*^*ε3/ε4*^ genotype increases risk for AD and related dementias. The new mouse model described here begins to address this and an *APOE*^*ε2*^ mouse line is currently being validated. However, further improvements may in mouse models could still be valuable to study APOE function. For instance, while the coding sequence of the mouse *Apoe* locus is humanized, the 5’ and most of the 3’ untranslated regions and regulatory elements are not. Some studies suggest important human-specific risk factors may reside within these non-humanized regions and humanizing additional portions of the mouse *Apoe* locus should be considered. However, it will be important to consider the binding of mouse proteins to human regulatory elements – a common issue when humanizing regions of the mouse genome.

Our results indicate that genotype plays an important role in cholesterol composition in the blood at 2 months, however this was not observed in a different cohort of mice at 4 months. With APOE being a key cholesterol trafficking protein, it was critical to characterize the lipid composition in the blood at the early ages studied. Cholesterol and lipids both contribute to fat homeostasis in the blood and the brain. However, cholesterol in mice is different to humans. Mice lack CETP which aids in the transfer of cholesterol esters depending on triglyceride concentration status (38). This inability to properly transfer cholesteryl esters and triglycerides has been shown to result in higher levels of HDL and lower levels of LDL (39). Further, it has been shown in humans that with each copy of *APOE*^*ε4*^, there is an increase in circulating triglyceride level (5, 40). Previous studies have shown that the *APOE*^*ε4*^ allele increases total plasma cholesterol and LDL concentration (41). While mice primarily use HDL, there were no differences between genotypes for HDL concentrations (41). Although human studies have reported differences in triglycerides due to *APOE* genotype, there was no evidence of an *APOE* genotype effect on triglyceride levels in our mice at 2 months. This may be due to the young timepoint studied in our experiment. Other studies using different human *APOE* targeted replacement mice, corroborated our findings showing that there was no significant difference in triglycerides between *APOE*^*ε3/ε3*^ and *APOE*^*ε4/ε4*^ mice at younger ages (42). At 4 months, although there was no overall effect of *APOE* genotype on cholesterol levels, we did observe a sex-activity effect on total cholesterol and HDL. While these findings indicate that *APOE* genotype plays a role on lipid homeostasis at an early age, aging studies in our mice are necessary to identify differential affects between *APOE* genotypes (e.g. *APOE*^*ε3/ε4*^ and *APOE*^*ε4/ε4*^) on cholesterol and triglyceride homeostasis that may lead to increased risk of dementia.

Transcriptional profiling of cortical tissue at 2 and 4 months revealed that the genes significant for *APOE*^*ε3/ε4*^ genotype enriched for different GO terms compared to the *APOE*^*ε4/ε4*^ genotype (using *APOE*^*ε3/ε3*^ as reference in our linear model). These data support our hypothesis that the *APOE*^*ε3/ε4*^ risk genotype confers unique effects compared to *APOE*^*ε4/ε4*^. The most significant term for *APOE*^*ε3/ε4*^ at 2 months included ‘sulfur compound binding’ which may indicate differences in homodimer formation. It has been shown that the APOE^ε3^ protein is able to form homodimers, as well as heterodimers with ApoA-II, which are believed to have different functions than the APOE^ε3^ monomers (43, 44). The APOE^ε4^ isoform however, cannot dimerize, as the critical cysteine is replaced by an arginine. These results signify the need to better understand the effect of the heterozygous state on the dimerization of APOE and how these differences may play a role in risk for dementia.

Linear modeling revealed several annexin genes were significant when comparing *APOE*^*ε3/ε4*^ to *APOE*^*ε3/ε3*^ but were not significant when comparing *APOE*^*ε4/ε4*^ to *APOE*^*ε3/ε3*^. Both *Anxa1* and *Anxa2* were differentially expressed and contributed to the functional enrichment terms ‘extracellular matrix’, ‘response to wound healing’, and ‘coagulation’. *Anxa2* has been shown to have direct effects on the plasminogen/fibrin clotting response (45). While we expected a linear increase in gene expression with increasing *APOE*^*ε4*^ gene dose, for endothelial-expressed *Anxa1, Anxa2*, and *Anxa3*, there was a significant increase in the *APOE*^*ε3/ε4*^ genotype when compared to *APOE*^*ε3/ε3*^, but not a significant increase in the *APOE*^*ε4/ε4*^ genotype when compared to *APOE*^*ε3/ε3*^. This pattern suggests that expression of some genes are differentially affected by the presence of both *APOE*^*ε3*^ and *APOE*^*ε4*^ alleles, which can help in our understanding of downstream effects of *APOE* ^*ε3/ε4*^ heterozygosity.

It is widely recognized that there is a female bias in *APOE*^*ε4*^-driven dementia risk, with postmenopausal females accounting for over 60% of AD cases (41, 46). In women, one *APOE*^*ε4*^ allele causes an increase in dementia risk equivalent to two *APOE*^*ε4/ε4*^ alleles in men (41). Our study supports that sex-specific *APOE*^*ε4*^ effects begin early in adolescence/adulthood and would be predicted to exacerbate dementia pathophysiology later in life. When examining the effect of sex-genotype for *APOE*^*ε4*^ status at 2 months and 4 months, there were more genes significant for sex-*APOE*^*ε4/ε4*^, than sex-*APOE*^*ε3/ε4*^. These data indicate that *APOE* genotype interacts with sex-specific characteristics at younger ages (even as early as adolescence) to differentially predispose females and males to dementia later in life.

Activity, particularly running, has been shown to have a protective effect on brain health in the context of dementias such as AD (47-49). Our experimental design incorporated this widely used intervention therapy to further extrapolate potential differences in *APOE*^*ε3/ε4*^ from *APOE*^*ε4/ε4*^ when compared to *APOE*^*ε3/ε3*^. While our study focuses on the importance of unique *APOE*^*ε3/ε4*^ transcriptomic signatures, we were also able to identify genes that were shared between the *APOE*^*ε3/ε4*^ and *APOE*^*ε4/ε4*^ genotypes. Linear modeling identified 50 genes that intersected between activity-*APOE*^*ε3/ε4*^ and activity-*APOE*^*ε4/ε4*^ that enriched for anoikis, which occurs when cells anchored to the ECM become detached. This suggests genes involved in ECM anchorage are similarly affected by running, regardless of the number of *APOE*^*ε4*^ alleles present. Further, to detect more subtle effects of *APOE* genotype on exercise, we performed WGCNA. While no modules enriched for both activity and *APOE* genotype, the cyan module correlated significantly with activity (**Supp Fig. 7**). The genes present in the cyan module enriched for myelination and gliogenesis. Previous studies have shown that voluntary exercise impacts myelination by increasing the proliferation oligodendrocyte precursor cells and mature oligodendrocytes in the motor cortex (50). It will be important to understand whether this may be protective or harmful in the long term. *APOE*^*ε4*^ typically exhibits harmful effects on the brain, predisposing to AD pathology and cognitive decline. In general, few studies have determined whether long-term exercise is beneficial in the context of *APOE* risk genotypes. Exercise may mitigate or exacerbate *APOE*^*ε4*^-dependent changes depending on the specific biological process considered.

Surprisingly, there were no significant functional enrichment terms for the genes unique to activity-*APOE*^*ε3/ε4*^ and activity-*APOE*^*ε4/ε4*^, although some patterns involving growth factors in activity-*APOE*^*ε3/ε4*^, and mitosis and protein folding in activity-*APOE*^*ε4/ε4*^, emerged from individual gene descriptions suggesting activity has subtle *APOE* genotype-specific effects in young mice that may become more pronounced with age.

## Conclusions

Due to *APOE*^*ε4/ε4*^ being less frequent in the human population, the *APOE*^*ε3/ε4*^ and *APOE*^*ε4/ε4*^ genotypes are usually concatenated in human studies, obfuscating certain characteristics that may be specific to the more common *APOE*^*ε3/ε4*^ genotype. Further, traditionally, mouse models have focused on comparing the *APOE*^*ε3/ε3*^ with *APOE*^*ε4/ε4*^. Our study predicts important differences between *APOE*^*ε3/ε4*^ and *APOE*^*ε4/ε4*^ genotypes on AD-relevant phenotypes including biometrics and cortical gene expression. These differences need to be better understood in order to properly determine whether the mechanisms increasing risk for diseases such as AD and related dementias in those carrying one *APOE* ^*ε4*^ allele are different from those carrying two particularly as differential *APOE* genotype effects may be exacerbated at older ages.

## List of Abbreviations

LOAD: Late Onset Alzheimer’s Disease
VaD: Vascular Dementia
HDR: Homology Directed Recombination
VLDL: Very Low Density Lipoprotein
LDL: Low Density Lipoprotein
HDL: High Density Lipoprotein
NEFA: Non Esterified Fatty Acids
WGCNA: Weighted Gene Co-expression Network Analysis
GO: Gene Ontology

## Declarations

### Ethics Approval and Consent to Participate

No human subjects or data was used in this study. All experiments involving mice were approved by the Animal Care and Use Committee at The Jackson Laboratory in accordance with guidelines set out in The Eighth Edition of the Guide for the Care and Use of Laboratory Animals. All euthanasia used methods were approved by the American Veterinary Medical Association.

### Consent for Publication

Not Applicable

### Availability of Data and Materials

All data is being uploaded to the AMP-AD Knowledge Portal. ID numbers will be provided once the process is complete.

### Author contributions

MS, GWC, GRH designed *APOE* mouse models. KEF and GRH conceived and designed this project. DTG, KPK, and KEF validated the *APOE* mouse models. KEF performed bioinformatic analysis. KEF, GWC, and GRH all consulted for statistical approach and analysis. KEF and GRH wrote and prepared this manuscript. All authors read and approved the final manuscript.

## Acknowledgements

Research reported in this publication was partially supported by the Jackson Laboratory’s Genetic Engineering Technologies Scientific Service. The authors wish to thank Todd Hoffert from Clinical Assessment Services for blood chemistry, Heidi Munger and the Genome Technologies group for RNA-sequencing, and Tim Stearns and Vivek Philip from Computational Sciences.

