## Supplementary figures and images for "*APOE*^ε3/ε4^ and *APOE*^ε4/ε4^ genotypes drive unique gene signatures in the cortex of young mice"

### Supplemental Figures

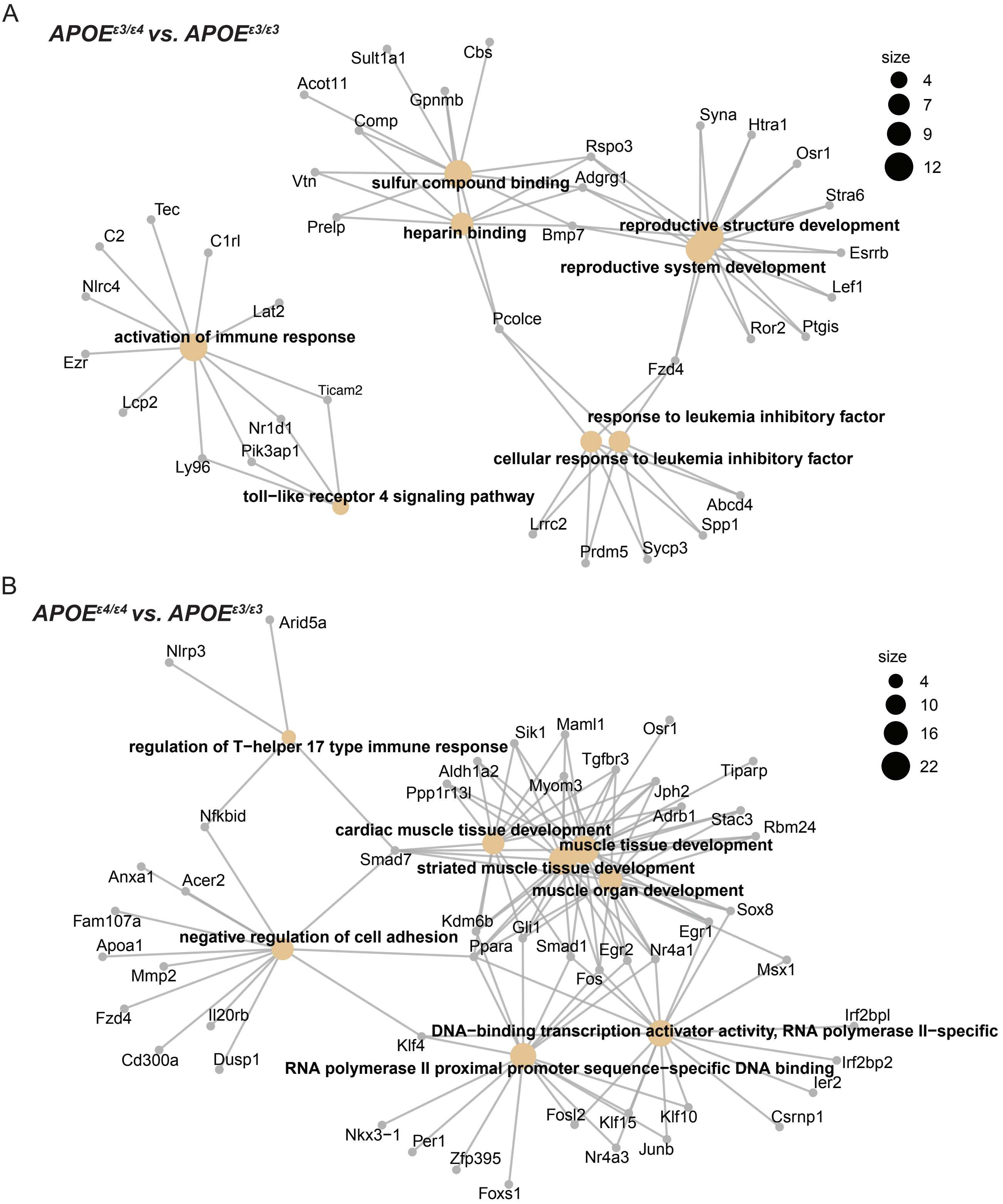

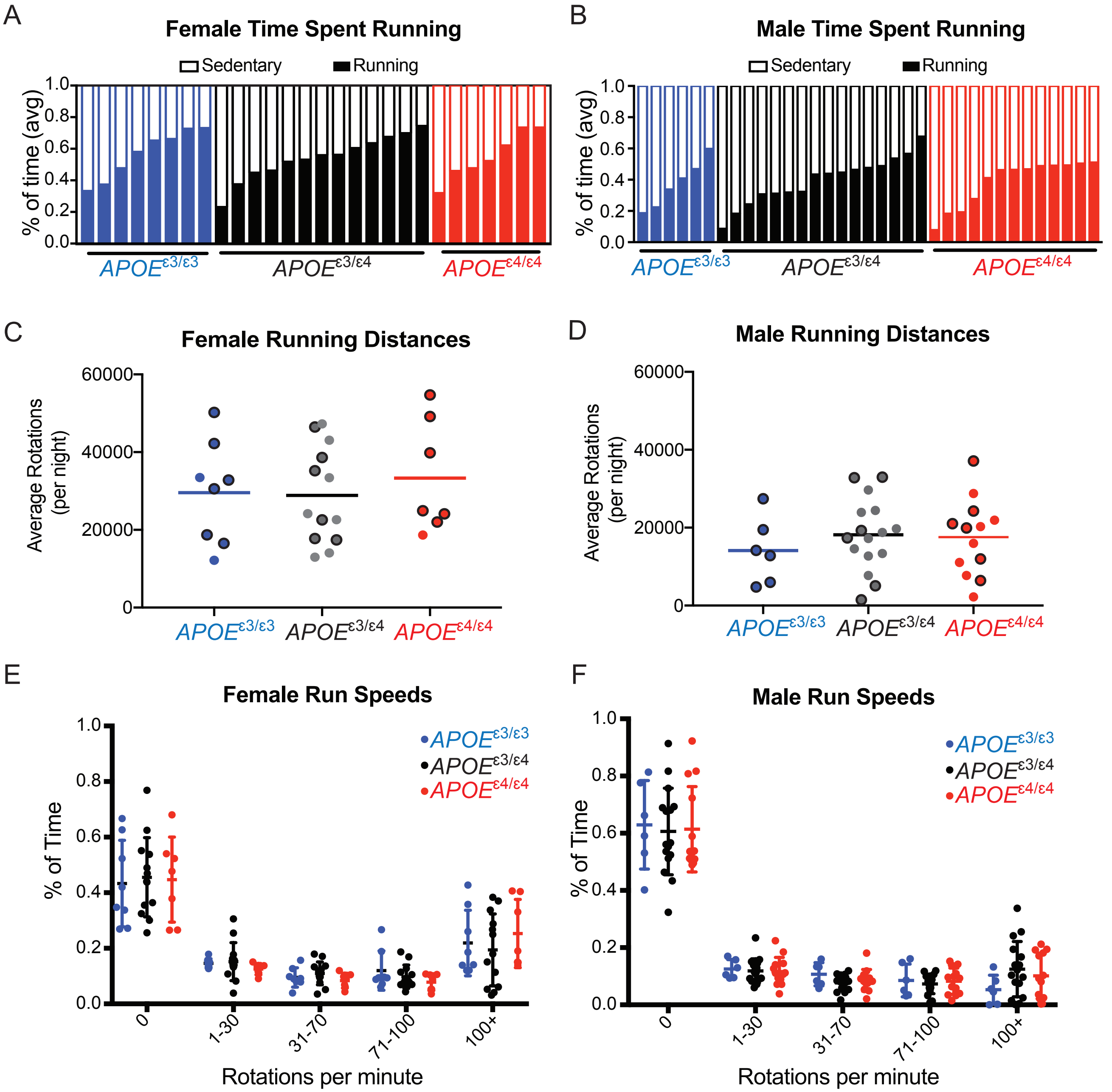

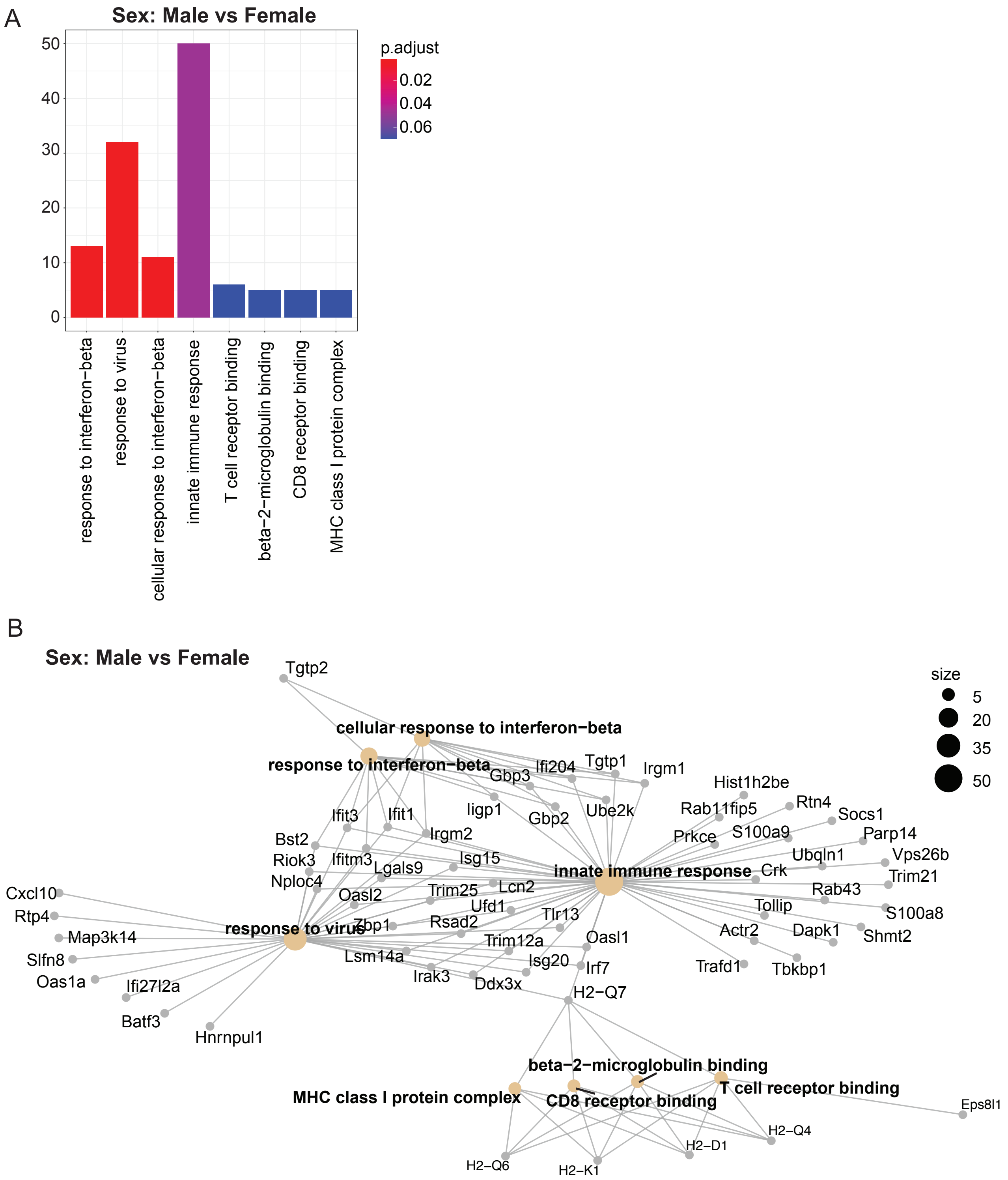

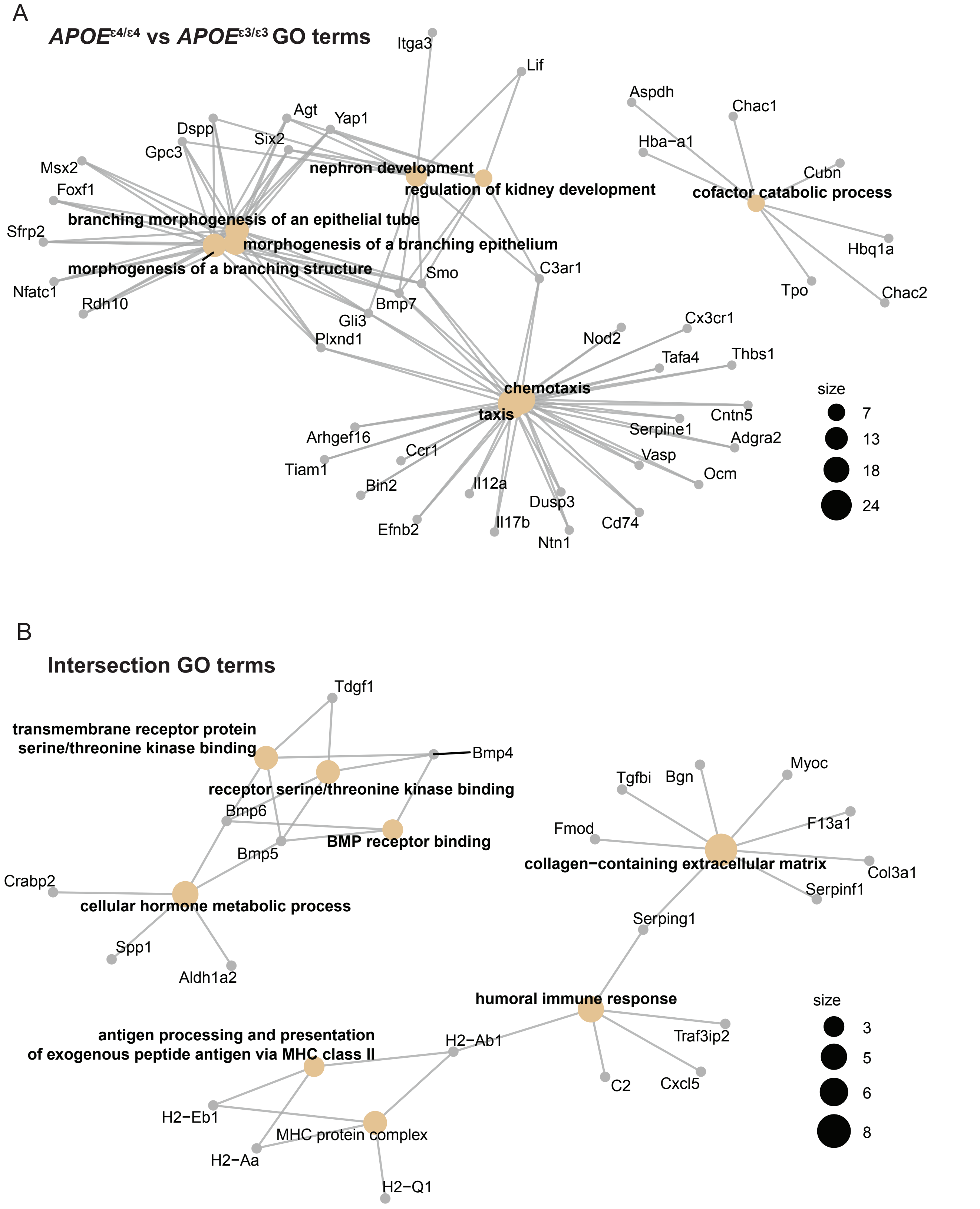

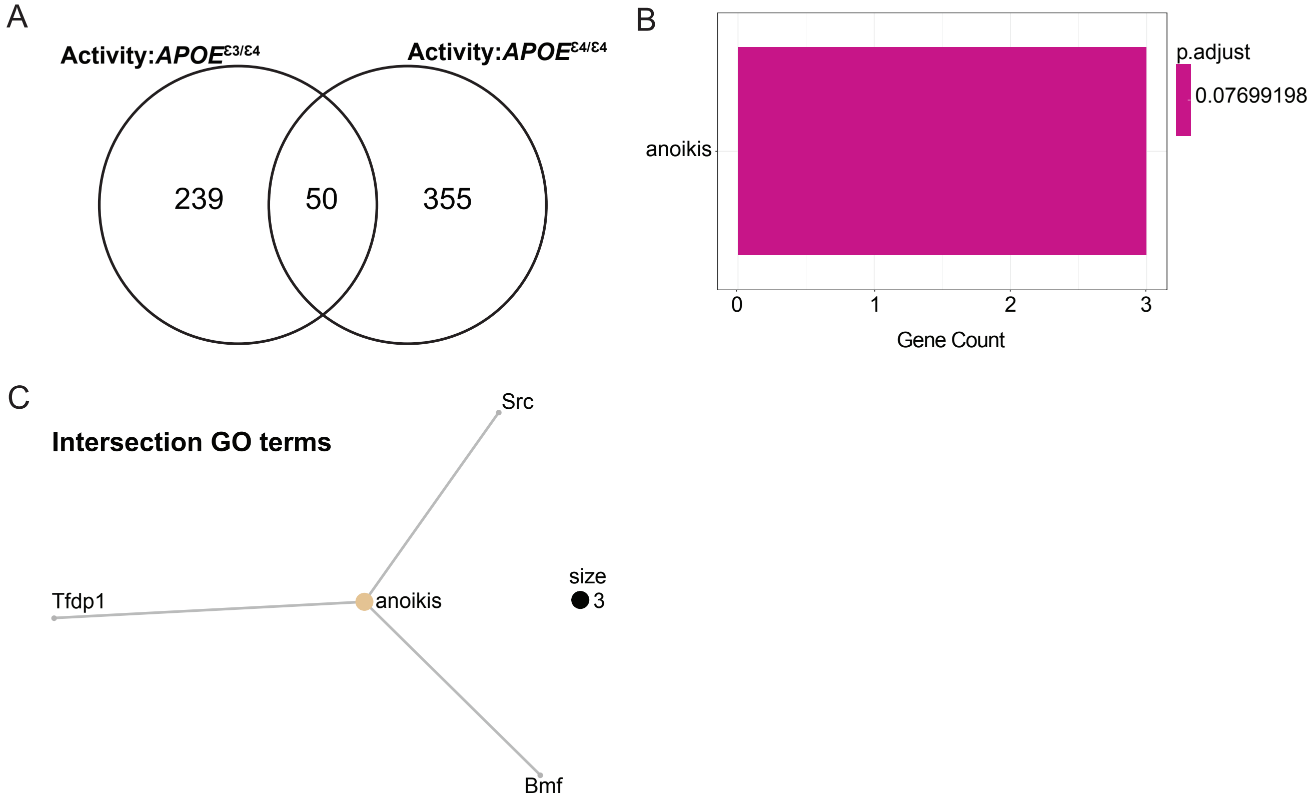

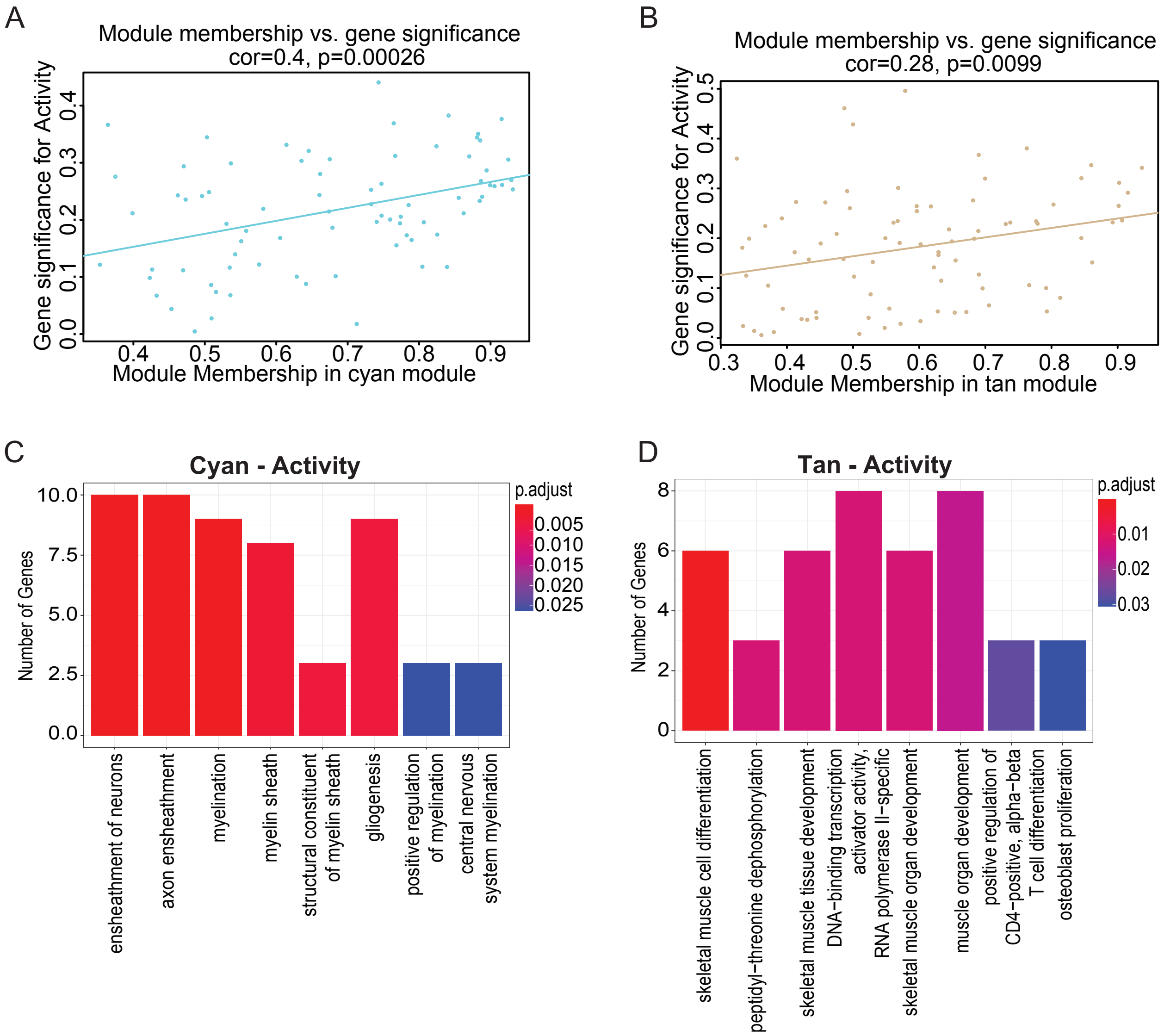

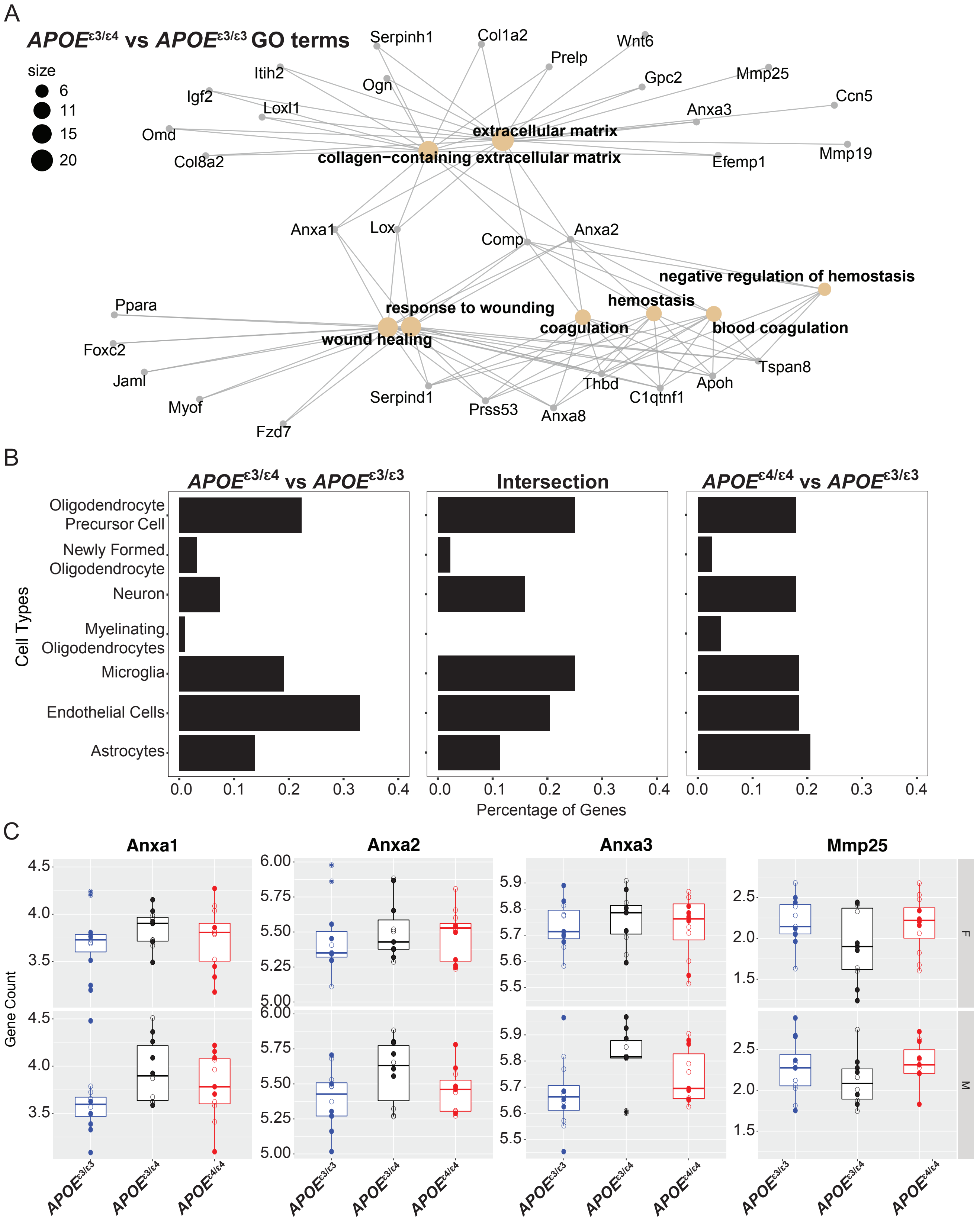

Mouse Apoe

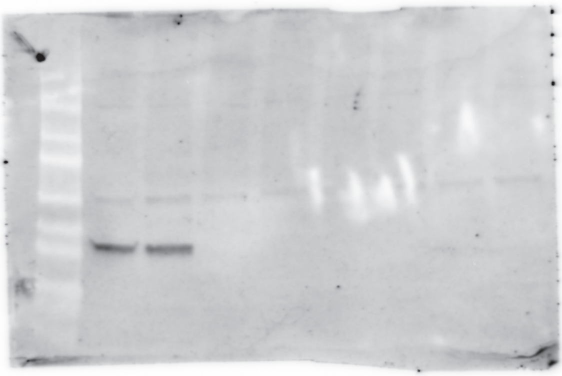

Human APOE

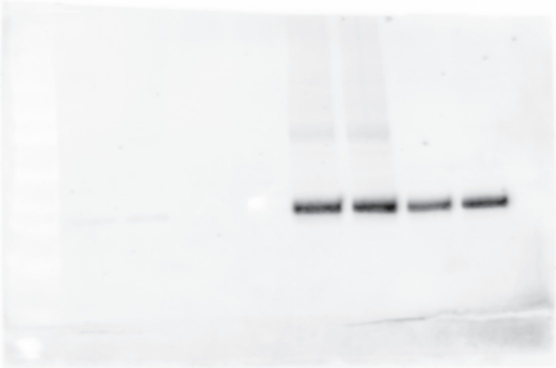

Human APOE$^{\varepsilon 4}$

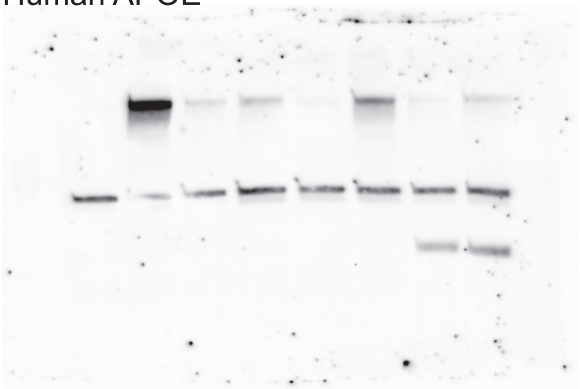

Actin

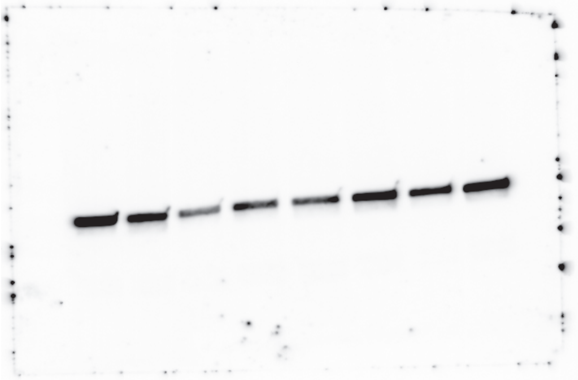
